# Adaptive evolution within the gut microbiome of individual people

**DOI:** 10.1101/208009

**Authors:** Shijie Zhao, Tami D. Lieberman, Mathilde Poyet, Sean M. Gibbons, Mathieu Groussin, Ramnik J. Xavier, Eric J. Alm

## Abstract

Individual bacterial lineages stably persist for years in the human gut microbiome^1–3^. However, the potential of these lineages to adapt during colonization of healthy people is not well understood^2,4^. Here, we assess evolution within individual microbiomes by sequencing the genomes of 602 *Bacteroides fragilis* isolates cultured from 12 healthy subjects. We find that *B. fragilis* within-subject populations contain substantial *de novo* nucleotide and mobile element diversity, which preserve years of within-person evolutionary history. This evolutionary history contains signatures of within-person adaptation to both subject-specific and common selective forces, including parallel mutations in sixteen genes. These sixteen genes are involved in cell-envelope biosynthesis and polysaccharide utilization, as well as yet under-characterized pathways. Notably, one of these genes has been shown to be critical for *B. fragilis* colonization in mice^5^, indicating that key genes have not already been optimized for survival *in vivo*. This lack of optimization, given historical signatures of purifying selection in these genes, suggests that varying selective forces with discordant solutions act upon *B. fragilis in vivo*. Remarkably, in one subject, two *B. fragilis* sublineages coexisted at a stable relative frequency over a 1.5-year period despite rapid adaptive dynamics within one of the sublineages. This stable coexistence suggests that competing selective forces can lead to *B. fragilis* niche-differentiation even within a single person. We conclude that *B. fragilis* adapts rapidly within the microbiomes of individual healthy people, providing a new route for the discovery of key genes in the microbiome and implications for microbiome stability and manipulation.

## Main Text

Billions of *de novo* mutations are generated daily within each person’s gut microbiome^6–9^ (**Table 1**). It is unknown if any of these mutations confer a strong adaptive benefit to the bacteria in which they emerge or, in contrast, all available mutations are deleterious or neutral. While some bacterial pathogens are known to adapt within individual infections^10–14^, investigations into healthy carriage of commensals have not revealed similar signals of within-person adaptive mutations^4,15^. These observations raise the possibility that millions of years of commensal evolution within mammalian digestive systems^16,17^ has exhausted all strongly beneficial point mutations. This hypothesis is echoed by signals of long-term purifying selection in the gut microbiome^2,18^. However, gut microbiomes are heterogeneous and individualized environments that may vary over time^1,14,19^, and it is possible that new mutations may still drive rapid adaptation of commensal species within individual people.

**Table 1:** Estimation of the number of mutations occurring daily within the human microbiome

| Number of bacteria per microbiome (cells/microbiome)<sup>5</sup> | Mutation rate of bacteria (SNP/nucleotide/replication)<sup>6</sup> | Average bacterial genome size (nucleotide/cell)<sup>7</sup> | Range of mean bacterial replication rate (replication/day)<sup>8</sup> | → | Estimated number of *de novo* mutation (SNP/microbiome/day) |
| --- | --- | --- | --- | --- | --- |
| $10^{13}$ - $10^{14}$ | $10^{-10}$ - $10^{-9}$ | $2\text{-}6 \times 10^6$ | 1-10 | | $2 \times 10^9$ - $6 \times 10^{12}$ |

Should adaptive mutations arise and be detectable within individual microbiomes, they are likely to indicate genes and pathways critical for long-term bacterial persistence in the human body^11,13,20,21^. The selective forces on these pathways might be common or person-specific, and their identification could guide microbiome-targeted therapies, including the selection and engineering of therapeutic bacteria for long-term colonization. To date, characterization of within-person evolution in the gut microbiome has been limited^1,2,4,22^, as it is difficult to distinguish *de novo* mutations from variants in homologous regions shared by co-colonizing bacteria using metagenomics alone. Culture-based approaches, which enable single-cell level whole-genome comparisons, have been limited to a small number of isolates. Further, it is often implicitly assumed that identifying within-person adaptation requires longitudinal sampling. However, if gut commensals diversify during their colonization within an individual, as is the case for bacterial pathogens^12,13,23^, co-existing genotypes can enable the inference of within-person evolution without long time-series.

To begin assessing the degree to which gut commensals evolve and diversify during colonization, we used a culture-dependent approach and focused on *Bacteroides fragilis*, a prevalent and abundant commensal in the large intestine of healthy people^24^. We surveyed intra-species diversity within 12 healthy subjects (ages 22-37; **Supplementary Table 1**), sequencing the genomes of 602 *B. fragilis* isolates from 30 fecal samples. These fecal samples included longitudinal samples from 7 subjects spanning up to 2 years and single samples from 5 subjects (**Supplementary Table 2**). None of these isolates were enterotoxigenic^25^ (Methods).

First using a reference based approach, we found that isolate genomes from different subjects differed by more than 10,000 single nucleotide polymorphisms (SNPs), while genomes from the same subject differed by fewer than 100 SNPs (with one isolate exception; **Extended Data Fig. 1**). We concluded that each subject was dominated by a unique lineage, consistent with previous investigations of within-host *B. fragilis* diversity^5,24,26^. We refer to each major lineage by its host ID (e.g. L01 for Subject 01’s lineage).

**Figure 1:**
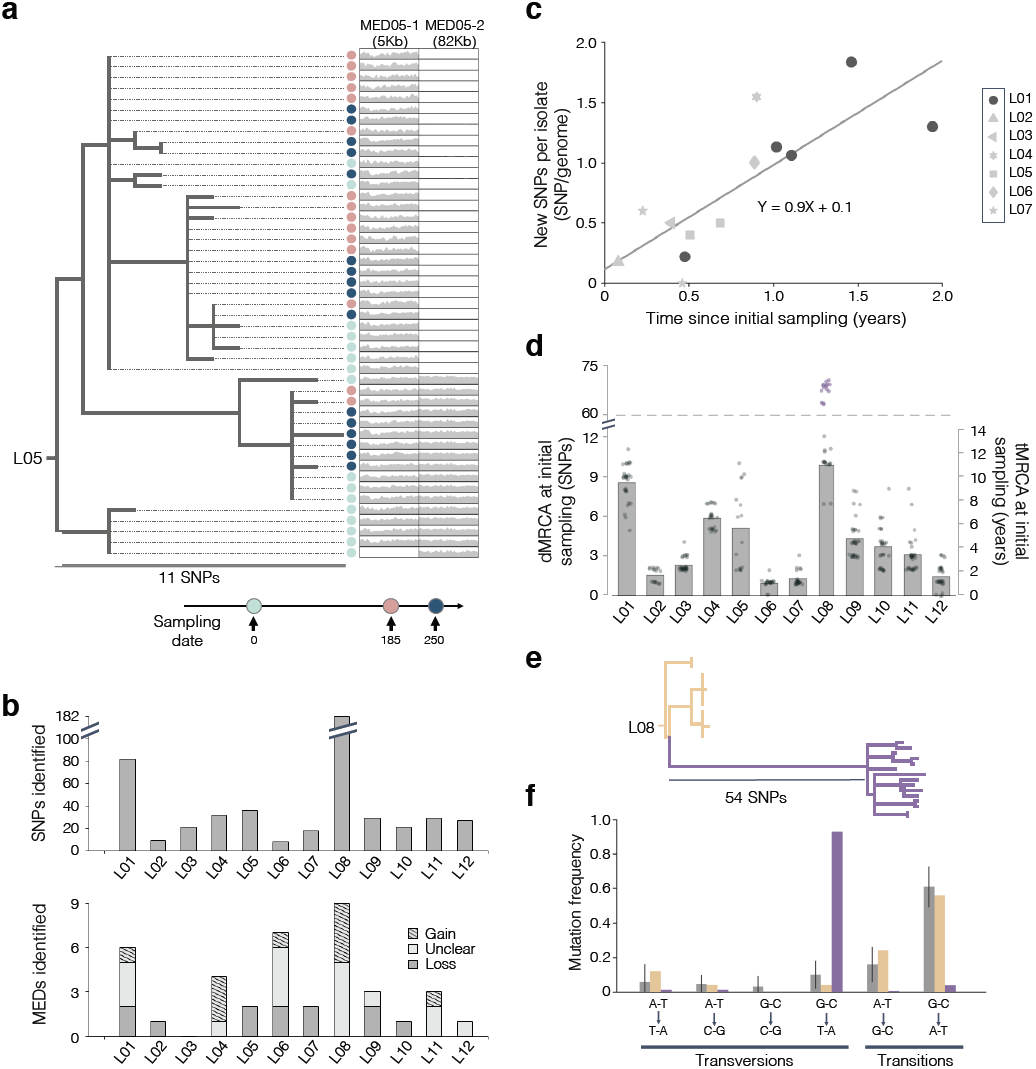
*B. fragilis* lineages diversify for years during colonization of healthy individuals via *de novo* SNPs and MEDs. (**A**) The phylogeny of isolates from L05 is shown as an example. Light blue, pink, and dark blue circles indicate isolates from samples taken at Day 0, 185, and 250, respectively. For each isolate, the relative coverage (compared to the mean genomewide) across the length of two mobile element differences (MEDs) is shown. For each isolate with an MED, the average relative coverage is ˜1X. (**b**) The number of SNPs and MEDs identified for each lineage. MEDs were classified as gained (hatched), lost (dark gray), or unclear (light gray). (**c**) Estimate of the molecular clock for *B. fragilis.* Each shape represents the average number of new SNPs per isolate not present in the set of SNPs at initial sampling of the same subject. Data points from L01 are colored in black, while data points from other lineages are colored in gray. (**d**) Bars represent average dMRCA and tMRCA at initial sampling for the 12 lineages. Gray dots represent individual isolates. For L08, the bar represents the mean for isolates from only the non-hypermutator sublineages and purple dots represent hypermutator isolates. (**e**) A rooted phylogeny of L08, with hypermutator sublineage (purple) and normal sublineages (yellow). The normal sublineages and the hypermutator sublineage share a single derived SNP. (**f**) The spectrum of mutations in the hypermutator sublineage (purple), normal sublineages of L08 (yellow), and 11 other lineages (gray). Gray bars represent averages across the 11 other lineages, and error bars represent standard deviation. See **Extended Data Fig. 5a-h** and Methods section for additional molecular clock and mutation spectrum analyses.

The SNP diversity was substantial within many lineages, enabling us to infer several years of within-person evolution. For each lineage, to discover variants in genomic regions not present in the reference, we assembled a draft genome using reads from all isolates, identified polymorphisms via alignment of short reads, and constructed a parsimony phylogeny (Methods, **Fig. 1a, Extended Data Fig. 2–4**). Between 8 and 182 *de novo* SNPs were identified per lineage (**Fig. 1b**). To estimate the age of the *B. fragilis* diversity within each subject at initial sampling, we calculated the average mutational distance of each population to its most recent common ancestor (dMRCA). To convert dMRCA to approximate units of time (tMRCA), we estimated the rate at which *B. fragilis* accumulates SNPs in the human gut by comparing SNP contents across longitudinal samples from the same subject (molecular clock; **Fig. 1c, Extended Data Fig. 5a-h**; Methods). Given our molecular clock estimate of ∼0.9 SNPs/genome/year, 11 of 12 lineages had values of tMRCA between ∼1.1-10 years (**Fig. 1d**) at the initial sampling. Due to the low acquisition rates of *Bacteroidetes* strains in healthy adult microbiomes^1,27,28^, we hypothesize that these within-subject populations emerged from a single cell within each subject. However, it is possible that some of the *B. fragilis* diversity within each person was inherited from a colonization event carrying multiple genotypes.

**Figure 2:**
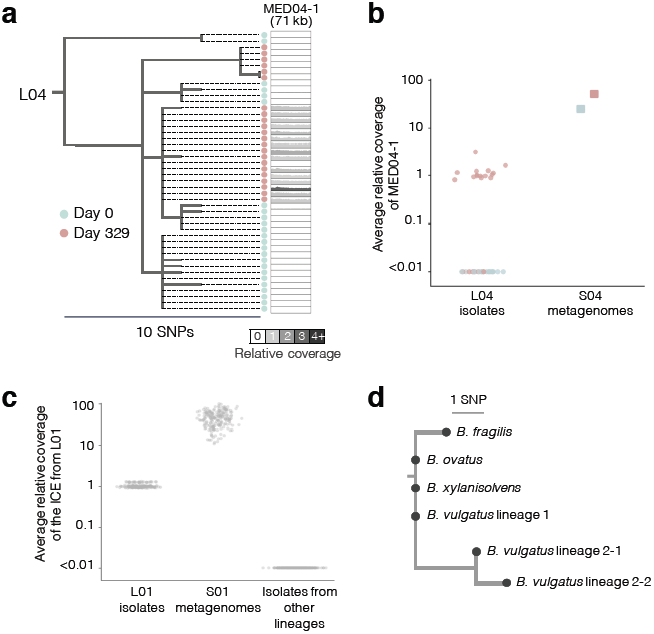
Mobile elements are transferred within the microbiome of individual subjects. (**A**) The phylogeny of isolates from L04, illustrating gain of MED04-1. Blue and pink circles indicate isolates taken at Day 0 and 329, respectively. Shading of the MED region reflects the average relative coverage of the MED in that isolate. (**b**) Average relative coverage across the length of MED04-1, a prophage, in L04 isolates (circle) and Subject 04 metagenomic samples (square). Colors represent sampling dates as indicated in (**a**). Isolates with this prophage had from 1X to 3X average coverage relative to the rest of the genome. (**c**) Average relative coverage of a putative integrative conjugative element (ICE) in isolates from L01, metagenomic samples from Subject 01, and isolates from other lineages. Isolates from the sample S01-0259 show slightly higher average relative coverage because genomic libraries of these isolates were prepared differently (Methods). (**d**) A rooted parsimonious phylogeny of the putative ICE across 4 species. Isolates that had identical ICE sequences and were from the same phylogenetic group are merged into a single node. In Subject 01, the *B. vulgatus* isolates were from 2 distinct lineages, one of which had 2 sublineages (**Extended Data Fig. 7**).

One outlier, L08, had a significantly higher value of dMRCA at initial sampling (38.9, P<0.001, Grubb’s test, **Fig. 1d**). This excess of mutations was due exclusively to an increase in a single type of mutation within one major sublineage (GC to TA transversions, P<0.001, Chi-square test), strongly suggesting a hypermutation phenotype (**Fig. 1e-f**, **Extended Data Fig. 5p**). Hypermutation, an accelerated mutation rate usually due to a defect in DNA repair, is associated with adaptation and its emergence is commonly observed in laboratory experiments and during pathogenic infections^23,29–32^.The dMRCA of non-hypermutator sublineages from L08 was compatible with within-person diversification (9.9 SNPs/genome/year), and the topology of the rooted phylogeny was also consistent with the emergence of a hypermutation phenotype within this subject (**Fig. 1e**). This is the first evidence of the co-existence of hypermutator and normal lineages within a healthy human, though commensal *E. coli* isolates with hypermutation phenotypes have been isolated before^33^.

Interestingly, each lineage’s tMRCA at initial sampling was less than its subject’s age (22-37 years), suggesting that these lineages colonized their subjects later in life, that adaptive or neutral sweeps purged diversity, or both. To determine if sweeps occur during colonization, we looked for mutations that fixed over time. We observed sweeps within 3 of the 7 lineages with longitudinal samples, one of which was associated with a significant decrease in dMRCA (L04; P <0.001; Wilcoxon rank-sum test; **Extended Data Fig. 5–6**). Thus, sweeps appear to be common during colonization, and *B. fragilis* lineages likely resided longer in their hosts than suggested by tMRCA at initial sampling.

We next assessed the contribution of horizontal evolution by identifying within-lineage mobile element differences (MEDs). We defined MEDs as DNA sequences with multi-modal coverage across isolates within a lineage (Methods). We found MEDs in 11 of the 12 lineages (**Fig. 1b**). These mobile elements include putative plasmids, integrative conjugative elements (ICEs), and prophages (**Supplementary Table 3**). We examined each MED’s distribution across its lineage’s phylogeny and used parsimony to categorize it as a gain or loss event. We inferred 10 MEDs gained, 12 lost, and 17 ambiguous loci in ∼50 cumulative years of evolution (using tMRCAs at initial samplings). This provided lower-bound estimates of ∼0.05 gain/genome/year and ∼0.04 loss/genome/year. We further estimated that MEDs change the *B. fragilis* genome by at least ∼1.3 kbp gain/genome/year and ∼1.9 kbp loss/genome/year. Thus, while gain and loss events are more rare than SNPs, they contribute more to nucleotide variation during *B. fragilis* evolution.

We reasoned that if these mobile elements were transferred from other species in the same microbiomes, we would observe evidence in metagenomes from the same stool communities. In particular, a transferred region should have increased coverage in the metagenome compared to the rest of the *B. fragilis* genome, owing to its presence in other species. We leveraged stool metagenomes available from 8 subjects, scanning for genomic regions with high relative coverage and high identity (>3X and >99.98%, respectively, Methods). We found evidence of one inter-species MED transfer within Subject 04 (38X relative coverage in the metagenomes; Methods; **Fig. 2a-b**). This MED, a putative prophage, was absent from all isolates at Day 0 yet present in 68% of isolates at Day 329. This combination of longitudinal genomic and metagenomic evidence strongly suggests that this prophage was acquired by *B. fragilis* during the sampling period.

This same approach enabled us to identify inter-species transfers even when the genomic regions were present in all *B. fragilis* isolates of a given lineage (for 3 lineages; **Supplementary Table 4**; **Fig. 2c**). We confirmed one candidate, a putative integrative conjugative element (ICE) in Subject 01 containing a type VI secretion system^34^ (T6SS), by culturing and sequencing 94 isolates of other *Bacteroides* species. This ICE was present in all isolates of 3 species (n=82) and contained only 4 SNPs among these species, suggesting recent transfer (**Fig. 2d, Extended Data Fig. 7,** Methods). T6SSs mediate inter-bacterial competition and have been shown to be shared by members of the same microbiome^24,35^. The prevalence of this ICE in this subject suggests it confers a strong selective advantage to its recipient species. In general, however, there are limited statistical tools for distinguishing adaptation from neutral evolution for mobile element changes.

To assess if positive selection was a significant driver of within-person *B. fragilis* evolution, we examined the identity of observed SNPs. We searched for parallel evolution, a hallmark of positive selection in which similar changes emerge independently, focusing specifically on parallel evolution occurring within a person. We identified 16 genes mutated in parallel within a single subject, a significant deviation from a neutral model (P<0.001, **Fig. 3a**, **Extended Data Fig. 8a-f**; Methods). These genes were significantly enriched for nonsynonymous mutations, as reflected by dN/dS, the normalized ratio of nonsynonymous to synonymous mutations, indicating that mutations in these genes were indeed adaptive (dN/dS = 6.03, CI = (1.57, 51.3)). In contrast, genes mutated only once within a subject did not show such an enrichment (dN/dS = 0.93, CI = (0.74, 1.16); **Fig. 3b**).

**Figure 3:**
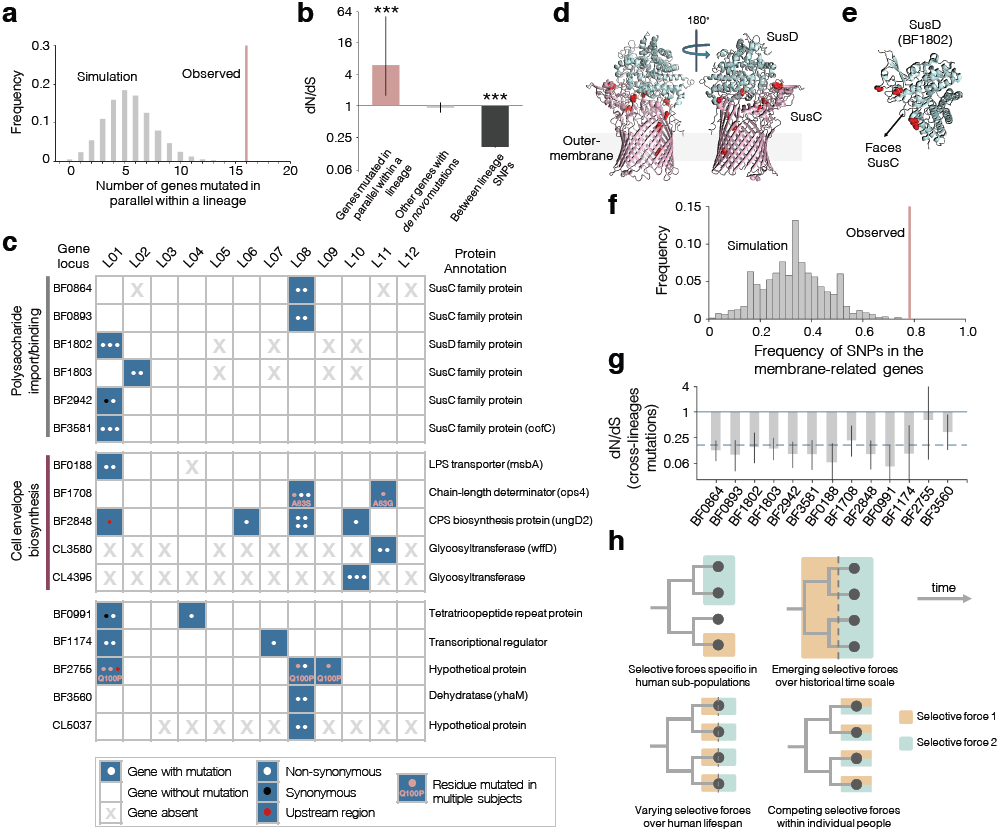
Genes involved in polysaccharide utilization and cell envelope biosynthesis are under selection to change during colonization. (**A**) Simulated under a null model (gray) and observed (pink) number of genes mutated in parallel within at least one lineage (P<0.001, Methods). (**b**) Canonical signal for selection, dN/dS, shows a signature of adaptive evolution for *de novo* mutations in genes under parallel evolution (P<0.001, Binomial test) but not for other genes. Mutations across lineages show a significant signature of purifying selection (P<0.001, Binomial test). Error bars represent 95% confidence intervals for dN/dS. (**c**) The 16 genes under parallel evolution are grouped by their biological functions and labeled with their locus tags and the inferred functions of their encoded proteins (**Supplementary Table 5**). Genes absent in a lineage are indicated with a gray X. Each dot in the table represents an independent mutation event, colored by type of mutation. (**d-e**) Mutations in SusC and SusD homologs under selection were enriched at the interface between the proteins (Methods). (**f**) Simulated under a null model (gray) and observed (pink) frequency of mutations in membrane-related genes (P<0.001, Methods). (**g)** Genes under parallel evolution show significant signatures of purifying selection across lineages (for 13 genes with inter-lineage mutations, Methods). Gray bars represent dN/dS for the gene, and error bars represent 95% confidence interval. The dashed line represents the average dN/dS for all inter-lineage SNPs. (**h**) Four models that could account for the discrepancy of natural selection at different timescales.

Genes under parallel evolution reveal challenges to *B. fragilis* survival *in vivo*. The 16 genes include 5 involved in cell envelope biosynthesis, a dehydratase implicated in amino-acid metabolism, and 4 with unclear biological roles (**Fig. 3c**). The remaining 6 genes all encode for homologs of SusC or SusD, a large group of outer-membrane polysaccharide importers (**Supplementary Table 5**). A typical *B. fragilis* lineage has 75 SusC/SusD pairs and their substrates are thought to be mainly complex yet unknown polysaccharides^36,37^. SusC proteins form homodimeric β-barrels capped with SusD lids^38^, and the observed mutations were enriched at the interface between the barrel and lid (**Fig. 3d-e**; Methods). Notably, one of these susC homologs (BF3581) has been shown to be critical for *B. fragilis* colonization in mice and its locus has been designated as commensal colonization factor (*ccf*)^5^. Its essentiality is thought to be related to binding to host-derived polysaccharides^5^, and, therefore, mutations altering Sus proteins might reflect pressures to utilize host or diet-derived polysaccharides^37^. Alternatively, the presence of Sus proteins in the outer membrane and their co-occurrence on this list with genes involved in cell envelope synthesis (**Fig. 3c, 3f**) hints that selection on these genes might be driven by the pressure to evade the immune system^39^ or phage predation^40^.

While our results show that single amino acid changes in key genes of *B. fragilis* confer rapid adaptive advantages within individual people, these same genes show signatures of purifying selection across lineages separated by thousands of years (**Fig. 3g**; Methods). Some of the mutated residues driving this adaptation are even highly conserved across species (>25% residues; **Supplementary Table 5**). The discrepancy in signals between timescales implies that the selective forces acting on these genes are not constant and raises the possibility that adaptive mutations occurring *in vivo* may incur collateral fitness costs in the context of other selective forces^41,42^. This notion of competing selective forces is echoed by the well-described invertible promoters of *B. fragilis,* which enable rapid alternation between different outer-membrane presentations^43,44^. Interestingly, the invertible promoters control the same major pathways that we identified as undergoing positive selection (capsule synthesis and polysaccharide importers)^43,45^. The non-constant selective forces driving these inversions and mutations might be specific to some people or lineages, recently introduced into the human population, present only at particular times (e.g. during early stages of colonization), or coexisting within individual people (**Fig. 3h**). We found evidence of both subject-specific and other selective forces. Three Sus genes (BF1802, BF1803, and BF3581) were each mutated multiple times within a subject, (P < 0.003 for each, Fisher’s exact test), yet no times in other subjects. In contrast, five genes under selection were mutated in multiple lineages, with two genes even acquiring mutations at the same amino-acid residue in different lineages (BF1708 and BF2755; **Fig. 3c**). Remarkably, a BF2755 mutation (Q100P) found polymorphic in 3 subjects was also in the ancestor of L12 and two publicly available genomes (**Extended Data Fig. 8g**), suggesting a common and strong selective pressure on this amino acid.

Could competing selective forces create multiple coexisting niches for *B. fragilis* even within a same individual? We noticed that the two lineages with the largest dMRCA at initial sampling (L01 and L08) had long-branched, co-existing sublineages that might reflect niche-differentiation (**Extended Data Fig. 2, 4a**). We closely examined L01’s evolutionary history over a 537-day period, during which the relative abundance of *B. fragilis* did not substantially change, using 206 stool metagenomes (**Extended Data Fig. 9a**). We tracked 21 abundant SNPs whose evolutionary relationships were previously identified from isolate genomes and inferred the population dynamics of their corresponding sublineages (**Fig. 4a-c**; Methods). The relative ratio of the two major sublineages (SLs), SL1 and SL2, which diverged ∼8 years prior to initial sampling, remained stable across the 1.5-year period (**Fig. 4c; Extended Data Fig. 9b**). SL1 showed multiple signatures of rapid adaptation during this period, including mutations in genes under selection, competition of mutations through clonal interference (e.g. between SL1-a and SL1-b, and within SL1-a), and a rapid sweep involving two SNPs related to Sus genes (0 to ∼70%<300 days; SL1-a-1; **Fig. 4c-d**). The continued coexistence of SL1 and SL2 despite a sweep within SL1 is particularly striking and suggests frequency-dependent selection or occupation of distinct, perhaps spatially segregated, niches^46–49^. The fact that 11 of 12 intragenic mutations separating these sublineages are amino-acid changing furthers the notion that they are functionally distinct. Therefore, it is likely that *B. fragilis* niche-differentiation can occur within a single person.

**Figure 4:**
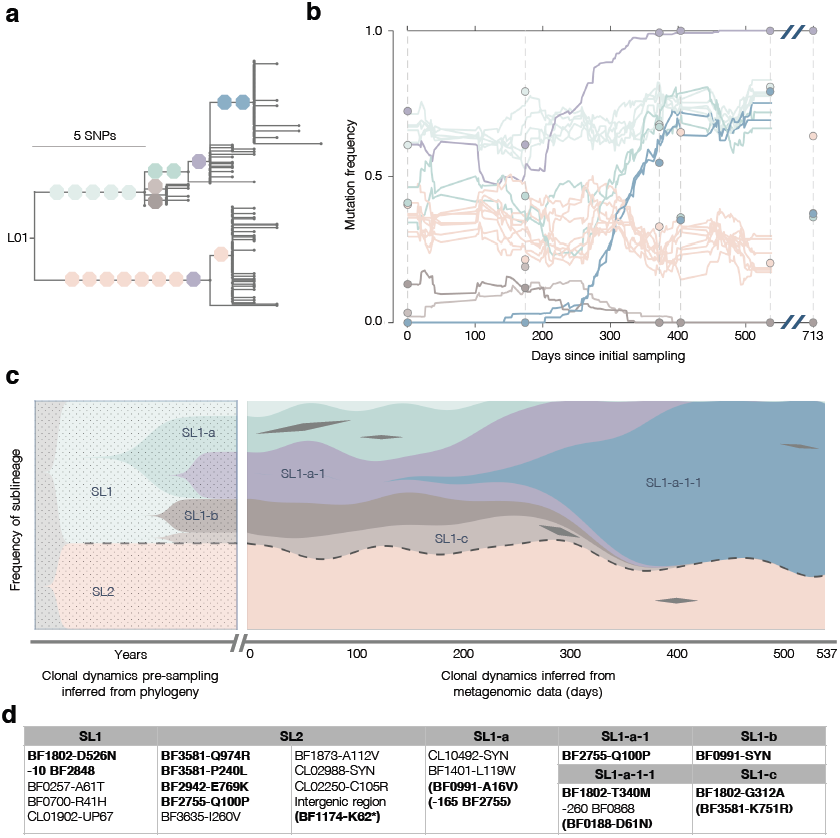
Two sublineages coexisted at a stable relative frequency despite rapid adaptive dynamics within one sublineage, suggesting niche differentiation within L01. (**A**) The phylogeny of isolates from L01. Branches with ≧4 isolates are labeled with colored octagons that represent individual SNPs. One SNP was inferred to have happened twice and is indicated in two locations (purple). (**b**) Frequencies of labeled SNPs over time in the *B. fragilis* population were inferred from 206 stool metagenomes (Methods). Colored circles represent SNP frequencies inferred from isolate genomes at particular time points. (**c)** The history of the sublineages carrying these SNPs prior to (left) and during (right) sampling was inferred (see Methods). The prior-to-sampling history is shaded to indicate temporal uncertainty. Sublineages are labeled with names and colored as in (**a**). Black diamonds represent transient SNPs from genes with multiple mutations. The two major sublineages, SL1 and SL2, are separated by dashed lines.(**d**) The identity of SNPs shown in (**c**) are listed in the table. SNPs in the 16 genes under positive selection are bolded and transient mutations in these genes are indicated with parentheses. Negative numbers indicate mutations upstream of the start of the gene. SYN indicates synonymous mutations.

We present here the first description of rapid within-person adaptation for a bacterial species whose native niche is the human intestine, as well as the most time-resolved description of bacterial within-person evolutionary dynamics to date. Within the gut microbiome of individual people, *B. fragilis* acquires adaptive point mutations in key genes, including polysaccharide importers and capsule synthesis genes, under the pressure of natural selection. This adaptation can be strikingly fast; near-daily tracking of one donor’s *B. fragilis* population revealed *de novo* mutations that rose from 0 to 70% frequency in less than a year (**Fig. 4b**). Continuing adaptation suggests there is no single optimal *B. fragilis* sequence for survival in the human microbiome and points to competing selective forces. Should rapid within-person adaptation be a common feature of gut commensals, as it is for many opportunistic pathogens of the cystic fibrosis lung^23,47,50^, it may have far-reaching implications for the microbiome field. Adaptation to the unique combination of selective forces present within each person may partially explain the observed stability of individual lineages in the microbiome^1^ and necessitate a personalized approach for microbiome manipulations. *De novo* mutation may need to be considered as a possible driver of ecological dynamics and inter-personal and temporal differences in community composition. Culture-based evolutionary approaches therefore provide both fundamental insights into the dynamics of human microbiomes and a powerful discovery route for genes and pathways critical to bacterial survival within the microbiome.

## Acknowledgements

We thank OpenBiome for providing stool samples, and Hera Vlamakis, Paige Swanson, Timothy Arthur, Julian Avila Pacheco, and Xiaofang Jiang for their assistance in obtaining samples and data. We are grateful to the BioMicroCenter at MIT and Microbial Omics Core at the Broad Institute for their assistance with library preparation and sequencing, Sean Kearney, Kathryn Kauffman, and Nadine Fornelos Martins for experimental assistance, and Vicki Mountain, Katya Frois-Moniz, and Shandrina Burns for administrative assistance. We thank members of the Alm lab for helpful discussions and Kevin Roelofs, Xiaoqian Yu, and Zhenrun Zhang for comments on the manuscript. This work was funded by a grant from the Broad Institute. T.D.L. acknowledges support from Boehringer Ingelheim.

## Author contributions

S.Z., T.D.L., and E.J.A. designed the study; S.Z. performed *B. fragilis* experiments; M.P. and M.G. performed experiments for other *Bacteroides*; S.M.G, R.J.X., and E.J.A. coordinated acquisition of metagenomic data. S.Z. and T.D.L. analyzed the data; S.Z., T.D.L., and E.J.A wrote the manuscript with input from all authors.

## Competing financial interests

Eric Alm is a co-founder and shareholder of Finch Therapeutics, a company that specializes in microbiome-targeted therapeutics.

## Methods

### Study cohort and sample collection

Stool samples were obtained from OpenBiome, a non-profit stool bank, under a protocol approved by the institutional review boards at MIT and the Broad Institute. All 12 subjects were healthy people screened by OpenBiome to minimize the potential for carrying pathogens and had ages between 22 and 37 years and body-mass indexes between 19.5 and 26.2 at initial sampling. Subjects were de-identified before receipt of samples. **Supplementary Table 1** contains detailed information about each subject.

OpenBiome received and processed fresh stool donations within 6 hours of generation. Most samples were homogenized in a buffer containing 12.5% glycerol and 0.9% sodium chloride by mass (relative ratio of buffer to stool was either 10:1 or 2.5:1 volume/mass). Some samples were homogenized in proprietary buffers (1:1 volume/mass). Homogenized samples were passed through a 330-micron filter and stored at −80°C. Subjects 01-07 had multiple samples from which *B. fragilis* was selectively cultured, with time-series spanning 31 to 709 days. For Subjects 08-12, only one sample was selectively cultured for *B. fragilis*. Metagenomic sequencing was performed on stool samples from 8 of the 12 subjects (319 stool samples in total). Detailed information about samples used for isolation, including handling conditions prior to sample receipt, is in **Supplementary Table 2** and information about samples used for metagenomic sequencing is in **Supplementary Table 6**.

### Library construction and Illumina sequencing

Samples were serially diluted in phosphate-buffered saline (PBS) and cultured for *B. fragilis* on *Bacterodies* Bile Esculin plates (BD 221836) in an anaerobic environment. Single colonies suspected of being *B. fragilis* based on colony morphology were re-suspended in 50μL of PBS with 0.1% L-cysteine. For future characterization, 15μL of the re-suspension was mixed with 15μL of 50% glycerol and stored at −80°C. DNA was extracted from the remaining 35μL using the PureLink *Pro* 96 genomic purification kit, following the manufacturer’s instructions. Genomic DNA libraries were constructed and barcoded using a modified version of the Illumina Nextera protocol^51^ (Library Prep. 1). Libraries from one sample (S01-0259, Day 709) were prepared by the BioMicroCenter at MIT using a different protocol, with lower input DNA and a final Pippin size-selection step (Library Prep. 2). Genomic libraries were sequenced either on the Illumina Hiseq platform with paired-end 100-bp reads or on the Illumina Nextseq platform with paired-end 75-bp reads by the Broad Institute Genomics Platform (**Supplementary Table 2**). Only isolates with average coverage of greater than 10 reads across the *B. fragilis* genome were included for analysis.

### Identification of major lineages and SNPs

To estimate the distance between isolates across subjects and identify major lineages, we aligned all short reads to a publicly available reference genome NCTC9343 (NCBI accession: CR626927.1) and identified SNPs. Reads were first trimmed and filtered using Cutadapt^52^ and Sickle^53^ (pe -f 20 -r 50), and aligned using Bowtie2 (Alignment parameters: -X 2000 --no-mixed --very-sensitive --n-ceil 0,0.01 --un-conc). Isolates for which more than 70% of reads aligned to the reference were included as being *B. fragilis*. From all subjects, 14 isolates were discarded (1 isolate from subject 10 and 13 isolates from subject 06), all of which had fewer than 5% of reads aligning to NCTC9343, suggesting other species. Candidate SNPs were identified using SAMtools^54^ and filtered using custom filters modified from previous work^23^. In particular, genomic positions were considered to be candidate SNP positions if at least one pair of isolates was discordant on the called base and both members of the pair had: FQ scores (produce by SAMtools; lower values indicate more agreement between reads) less than −60, at least 7 reads that aligned to each of the forward strand and reverse strand, and a major allele frequency of at least 90%. If the median coverage across samples at a candidate position was less than 10 reads or if 33% or more of the isolates failed to meet filters described above, this position was discarded. For each SNP position identified, a nucleotide call was assigned to each isolate using the major allele call across reads for that isolate at that position. If fewer than 7 reads aligned to either forward or reverse strand of a position in an isolate, or the major allele frequency was smaller than 90%, an ambiguous call was assigned to the isolate at that SNP position.

We generated a neighbor-joining tree from the concatenated list of variable positions from conserved genomic regions present in all *B. fragilis* isolates from all subjects. When computing the distance between each pair of isolates, we only used variable positions that had unambiguous nucleotide calls from both isolates. This tree showed 12 major clades corresponding to the 12 subjects and one minor clade containing a single isolate from Subject 10 (**Extended Data Fig. 1a**). Within each major clade, all isolates differed from one another by fewer than 100 SNPs. We therefore operationally defined a lineage as a set of isolates that differ by fewer than 100 SNPs and refer to specific genotypes within a lineage as sublineages. All lineages differed by over 10,000 mutations (**Extended Data Fig. 1b**); given the molecular clock estimated by this work, this represents at least thousands of years of evolutionary distance.

### *De novo* assemblies of lineage genomes and within-lineage SNP identification

To enable us both to detect variants within genes carried only in a subset of lineages and to detect gains and losses of genomic regions that are specific to single lineages, we created a pan-genome for each major lineage. For each major lineage, we concatenated reads (trimmed and filtered) from all isolates and used this concatenated file as the input for *de novo* genome assembly via Spades v3.10.0 (parameter: --careful)^55^. To limit the memory required for assembly, we used 0.25 million pairs of reads from each isolate (∼7x coverage). Isolates prepared by the Library Prep. 2, as well as a few isolates with apparent cross contamination (genome assemblies built only using reads from single isolates were larger than 6MB; *B. fragilis* genome assembly sizes range from 4.8 to 5.3 MB) were excluded in building assemblies. Isolates not used to build the genome assembly are indicated as such in the metadata associated with the uploaded raw data (see **Data availability**). Statistics of these genome assemblies are in **Supplementary Table 1**.Assembly genomes were annotated using Prokka v1.11^56^. Lineage pan-genomes successfully assembled regions present in only a single isolate (e.g. **Extended Data Figure 2, 3c, 3e**) and enabled detection of mutations that would have been missed by comparison to a single reference (**Extended Data Fig. 3c v**s **Extended Data Fig. 3g**). A genome assembly of the minor lineage from Subject 10 was built using all reads from this isolate.

Within-lineage mutations were identified by alignment of short reads to the corresponding lineage genome assembly, using the same parameters as described in the previous section. For lineage 10, the major allele frequency filter was set to 95%. Candidate positions in MEDs were also discarded (see below for information on MED identification). Detailed information of intra-subject SNPs from the 12 subjects are listed in **Supplementary Tables 7-18**.

The gene content across the 12 major lineage genomes and the NCTC9343 reference varied between 10%-20% (Using the Szymkiewicz-Simpson similarity coefficient and taking gene length into account, **Supplementary Table 19**).

### Toxin detection

We compared the genome assemblies of the 12 major lineages and 1 minor lineage to the Virulence Factors Database, which contains >2400 virulence factors^25^, via BLAST using a threshold bit score of 200. We found only two hits to the database: Cps4J in L11 and ospC4 in L01. Both hits were not toxins previously characterized for *B. fragilis*. In contrast, this method identified 171 hits to known *B. fragilis-*related toxins from 30 out of 88 *B. fragilis* genomes from National Center for Biotechnology Information (NCBI).

### Phylogeny of isolates from each *B. fragilis* lineage and identification of ancestral alleles

We used parsimony to reconstruct the evolutionary relationship between isolates from the same lineage. For each major lineage, a phylogeny of all isolates was built using a list of concatenated intra-subject SNPs and the closest lineage as an outgroup. We used the dnapars program, a parsimony tree builder from PHYLIP v3.69 to infer the phylogeny^57^. When parsimony could not resolve which allele was more likely to be ancestral, we inferred the ancestral allele to be the majority nucleotide at this genomic position across all other lineages with this genomic region. If a region was unique to a lineage, we assigned the ancestral allele that minimized the average mutational distances to the most recent common ancestor (dMRCA) for all isolates (3 cases).

### dMRCA of each *B. fragilis* major lineage, molecular clock, and tMRCA

To calculate dMRCA for each subject at each time point, we counted the number of positions at which the called allele was different than the ancestral allele for each isolate, assessing only SNP positions that were polymorphic among isolates from the particular time point, and averaged the results.

For each lineage with multiple time points, we computed the average number of new SNPs brought in per isolate from a later time point compared to the collection of SNPs identified at the initial time point. We then used linear regression to estimate the rate of evolution. The slope of the regression is our estimation of the evolutionary rate (**Fig. 1c**). Additional analysis approaches gave similar values of the molecular clock (**Extended Fig. 5a-h**).

Each tMRCA was calculated by dividing dMRCA by the estimated molecular clock (**Fig. 1d**). We stress that tMRCA is not an estimate of time to colonization, but simply an estimate of the age of the coexisting diversity, as sweeps can purge diversity. While potential systematic false negative and false positive SNPs may have impacted tMRCA values, these sources of error would have had a similar impact on our molecular clock estimation, as SNP-calling was consistent throughout. Other possible sources of error in estimating tMRCA include incorrect designation of ancestral versus derived allele and undersampling of the population, though collector curves for dMRCA indicate that sampling was usually sufficient (**Extended Data Fig. 6a-l**). Interestingly, collector curves for the number of *de novo* SNPs reflect that the number of SNPs identified did not saturate (**Extended Data Fig. 6m-x**).

### Mutation spectrum of hypermutator sublineage

SNPs were categorized into 6 types, based on the chemical nature of the single nucleotide changes (**Fig 1f**). For L08, we computed the frequency of each type separately for the hypermutator sublineage and non-hypermutator sublineages (**Fig. 1f**, purple and yellow bars). For the remaining lineages (L01-L07 and L09-L12), we computed the mutation spectrum for each lineage and then computed the mean and standard deviation of each of the 6 types (**Fig. 1f**, gray bars). The mutation spectrum was significantly different between the hypermutator sublineage and the non-hypermutator sublineages (Chi-square test, P<0.001), as well as the mean across the other 11 lineages (Chi-square test, P<0.001). No significant difference was found between the 11 other lineages and the non-hypermutator sublineages from L08 (Chi-square test, P=0.4).

When excluding the GC-TA type of mutation from the analysis, we found no significant difference between the non-hypermutator sublineage in L08 from the 11 other lineages (**Extended Data Fig. 5p**, P=0.11, Chi-square test), suggesting that the hypermutation phenotype was exclusively due to an increase in GC-TA mutations.

### Identification of Mobile element differences (MEDs)

We aligned short reads to the assembled genome of each major lineage as above and identified candidate regions that were at least 500nt in length, had low relative coverage (< 0.2X) at every nucleotide in at least one isolate, and had >0.9X coverage at every nucleotide in at least one isolate. For L01, we excluded isolates from the last time point, as these isolates’ genomic libraries were prepared differently than the other isolates and therefore had different coverage pattern genomewide.

To account for the fact that single mobile elements could have been separated into multiple pieces in the genome assembly, we grouped regions suspected to emerge from the same event. We clustered sequences that had identical presence/absence patterns across all isolates, where presence was defined by >0.4X average relative coverage over the region. On 3 occasions, we noticed regions that had the same presence/absence pattern but had different coverage distribution across isolates, suggesting they came from distinct mobile elements. In these cases, we separated these clusters of sequence regions into clusters with consistent coverage distribution patterns. Detailed information of all MEDs is in **Supplementary Table 3**.

### MED gain and loss rates

We used parsimony to infer whether a MED was a gain or loss event. For each MED, we inferred events on the phylogenetic tree generated from whole genome data. If a single change of one type (e.g. gain) could explain the distribution, but more events were required for the other type (e.g. loss), the MED was categorized as such (**Supplementary Table 3; Fig. 1b**). Seventeen MEDs were classified as unknown because either: multiple gain or multiple loss events were required to explain the distribution (e.g. MED01-2); or both a single gain event and a single loss event were consistent with the distribution. Interestingly, one putative MED from L11 appeared to have been lost many times among isolates during culture (**Extended Data Fig. 4d, f**). To estimate lower bounds for the rates at which gain and loss events change *B. fragilis* genomes, we weighted each observed MED *j* by its frequency within lineage *i* (*f*_*ij*_). We then divided the weighted sum of events by the total time of diversification, estimated by the sum of tMRCA at initial sampling. The following equation was used for gain and loss events, separately:

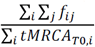

To estimate the absolute contribution of gain and loss events to the size of *B. fragilis* genomes, we accounted for length of each MED (*L*_*ij*_).

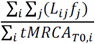

### Metagenomic library construction and Illumina sequencing

Genomic DNA was extracted from stool samples for metagenomic sequencing by the Microbial Omics Core at the Broad Institute using MoBio PowerSoil kits (Qiagen 12955-4) according the manufacturer’s instructions. Genomic DNA libraries were constructed and barcoded by the Broad Technology Labs from 100-250pg of DNA using the Nextera XT DNA Library Preparation kit (Illumina) according to the manufacturer’s recommended protocol, with reaction volumes scaled accordingly. Pooled libraries were sequenced on the HiSeq platform with paired-end 100bp reads by the Broad Technology Labs.

### Inter-species mobile element transfer

For each lineage, we scanned the assembled genome for regions with high average relative coverage when aligning metagenomic reads to the lineage genome assembly (>3X). The coverage of metagenomic reads over the *B. fragilis* assembly varied over as much as 1000 folds due to reads from homologous regions of different species. Therefore, to normalize against the true expected coverage of the *B. fragilis* genome, we divided observed coverage at each position by the mean coverage across positions between the 30^th^ percentile and 70^th^ percentiles (median was not precise given the low coverage in some samples). To identify recent transfer events, we searched the genome for candidate regions >5000 nucleotides in length and in which the consensus genome from metagenomes was <0.02% different from the consensus genome from isolates of the same subject. We found 14 candidate regions in 3 lineages. We found only two candidate regions that overlapped with MEDs, all of which were in Subject 04 (representing one MED). Information about these candidate regions is listed in **Supplementary Table 4**.

We identified two genomic regions (31 Kb and 62 Kb, respectively) that were candidates for inter-species mobile element transfer in Subject 01. These two regions contained distinct ORFs homologous to conserved genes from type 6 secretion system of genomic architecture 2 (**Extended Data Fig. 7c**), consistent with a single transfer event. This transfer event was inferred to be an integrative conjugative element (ICE) because it contains the *tra* genes associated with integrative conjugative elements and a tRNA gene at one edge of a transfer region (**Supplementary Table 4**). To test if the putative ICE was indeed transferred between species, we cultured and sequenced the genomes of 94 *Bacteroides* isolates from this subject. We examined 53 *Bacteroides vulgatus* isolates (43 isolates one *B. vulgatus* lineage, 10 isolates from a different *B. vulgatus* lineage, **Extended Data Fig. 7a, b**), 25 *Bacteroides ovatus* isolates, 4 *Bacteroides xylanisolyens* isolates, 10 *Bacteroides stercoris* isolates and 2 *Bacteroides salyersiae* isolates. We sequenced these isolates as described for *B. fragilis* and aligned reads to the mobile element candidates, using the same parameters for *B. fragilis.* Strikingly, both genomic regions were present (average coverage >10 reads) in all *B. ovatus, B. xylanisolyens,* and *B. vulgatus* isolates profiled, but absent in all isolates of the other two species. The perfect co-occurrence of these two genomic regions further supports that they were from a single transfer event.

### Parallel evolution

We counted a gene as under parallel evolution if, in at least one subject, the gene had multiple independent SNPs and more than 1 SNP per 2,000 bp (to account for the fact that long genes are more likely to be mutated multiple times by chance). Cases in which two SNPs in the same gene always occurred together in the same isolates were not included as parallel evolution (one case from L04). To identify nucleotide positions that mutated multiple independent times within a person, we leveraged the parsimony phylogenies described above. We inferred the genotypes of all internal nodes using the parsimony assumption and counted the number of mutation events. This method identified 3 nucleotides that were mutated multiple times within an individual **(Extended Data Fig 2, 4)**. To determine whether the number of genes under parallel evolution represented a significant departure from what would be expected in a neutral model, we performed for each subject 1,000 simulations in which we randomly shuffled the mutations found across the lineage genome assembly and calculated how many genes showed a signature of within-person parallel evolution (**Fig. 3a**). To compare genes from different assemblies, coding sequences identified by Prokka from all lineages were clustered using CD-HIT with at least 98% identity and 90% coverage^58^. Detailed information for each gene under parallel evolution is in **Supplementary Table 5** and gene clusters are listed in **Supplementary Table 19.** Simulations performed for metrics of cross-subject parallel evolution did not yield additional signatures of adaptive evolution (**Extended Data Fig. 8**).

#### dN/dS

Mutations were categorized as synonymous (S) or non-synonymous (N) based on open-reading frame annotations created by Prokka^56^. To calculate dN/dS for sets of *de novo* mutations emerged within subjects (**Fig. 3b**, first two categories), we normalized the observed N/S ratios by the expected N/S ratios. For any given set of SNPs, we calculated the expected N/S for these SNPs, accounting for both (1) the different probabilities of acquiring nonsynonymous mutations for different types of mutations and (2) the codon compositions of the genes in which these SNPs occurred. This method is similar to what we have done previously^23^, but accounts for different codon composition between genes. 95% confidence intervals were calculated using binomial sampling.

To compute dN/dS for mutations across lineages (**Fig. 3b**, third category), we leveraged publicly available sequences. We downloaded fastq files of 55 publicly available *B. fragilis* isolate sequencing runs. We then identified mutations across these genomes and the 12 major lineages from this study (one isolate per lineage) using the same approach and parameters described above (Identification of major lineages and SNPs). The NCTC9343 genome was used as reference and ancestor. Expected N/S ratio was calculated with the same method described above, using all the SNPs identified across lineages.

To compute dN/dS for cross-lineage mutations in individual genes (**Fig. 3g**), we normalized the observed N/S with expected N/S of the particular genes. Expected N/S ratio was calculated with the same method described above, using only cross-lineage SNPs identified within the particular genes. For 3 genes not present in the NTCT9343 genome, we used the *de novo* assemblies to recruit reads from the publicly available sequences. No cross-lineage SNPs were identified in these 3 genes and dN/dS was not reported for these genes.

### Annotation of genes under selection

To discover homologs of the sixteen genes under within-person parallel evolution, we used blastp to search against the RefSeq database, excluding proteins from *B. fragilis* genomes. Top hits with 3-4 letter gene names were searched against the *B. fragilis* genome to confirm whether they are true orthologs. We used the organisms from which these gene names were initially described to avoid false propagation of misannotation. We also used PaperBLAST to aid in identifying candidate gene names^59^. Cellular localizations were predicted using CELLO.

Conservation scores for each mutated residue was predicted using the Consurf web service^60^. For each gene, we used blastp to find homologs from the RefSeq database (first 100 hits; sequence similarity from 35% to 95%; query coverage > 80%). A multiple sequence alignment (MSA) was created using Clustal omega from the EMBL-EBI web service {ref} (default parameters). We then used each MSA to generate conservation score at each amino-acid residue using Consurf (default parameters). Detailed information is in **Supplementary Table 5.**

### SusC and SusD protein structures and interface residues

Available crystal structures of a SusC homolog (BT1763) from *Bacteroides thetaiotaomicron*^38^ and BF1802 from *B. fragilis* NCTC_9343^61^ were used to visualize the mutations observed in Sus genes under parallel evolution. We aligned the five *B. fragilis* SusC proteins under parallel evolution and BT1763 using Clustal Omega from the EMBL-EBI web service^62^ (default parameters). For all non-synomymous mutations, we identified their aligned positions on the BT1763 crystal structure. Two amino acid residues aligned to the first 211 amino-acid region, which encodes for a plug domain and is not available in the crystal structure of BT1763^38^. Eight non-synonymous mutations from Sus genes under parallel evolution are marked in red in **Fig. 3d** and **Fig. 3e**, using PyMol software^63^.

To test if the mutated residues were enriched at the interface between SusC and SusD, we used the PDBePISA web service^64^ (default parameters) to classify residues on the BT1763 crystal structure as in contact or not in contact with the SusD homolog. Of 806 residues, 119 were inferred to be interface residues. Among the 8 residues that were mutated in parallel, 4 of them were predicted to be interface residues in both programs, a significant enrichment (P=0.02, Fisher exact test). A similar result was obtained using the PyMol function InterfaceResidues (cutoff=1.0; P=0.02, Fisher exact test).

### Enrichment of membrane proteins

For all genes from the 12 major lineage genome assemblies, we used CELLO^65^ to predict the cellular localization. Genes were considered to be membrane-related if they were annotated as inner membrane, periplasmic, or outer membrane. To compare our observation to the null expectation, we performed simulations. For each of the sixteen genes, we randomly selected one gene from the genome assembly of the lineage in which parallel evolution was identified. If a gene had parallel mutation in multiple lineages, we randomly chose one of the lineages. The cellular localization of *n* SNPs was assigned based on the CELLO prediction of this randomly picked gene, where *n* is the number of SNPs the original gene had across lineages. The proportion of SNPs from membrane-related genes was inferred using all sixteen such randomly picked genes (repeat genes not allowed). This procedure was repeated 1000 times to draw a null distribution of proportion of membrane-related SNPs. We calculated that in the sixteen genes under selection, 79% of the SNPs are from membrane-related genes, a significant deviation from the null distribution (P<0.001, **Fig. 3f**).

### Signatures of subject-specific adaptation

Fisher’s exact statistic was used to test subject-specific adaptation, comparing the number of SNPs in a tested gene within a particular lineage, the number of SNPs in other genes within this lineage, the number of SNPs in this gene from all other lineages combined, and the number of SNPs in other genes from all other lineages combined. We tested 10 genes that were present in multiple subjects but mutated only in one subject. The p-values for BF1802, BF3581, BF1803, are all less than 0.005, suggesting person-specific adaptation.

### Mutation dynamics

Metagenomic reads from Subject 01, acquired as described above, were aligned to the assembled genome of L01 using the same parameters described for aligning isolates reads. We tracked the frequency of each SNP found in 4 or more isolates from L01; SNPs found in fewer isolates were not abundant in the metagenomes. For each of the 21 SNPs that met this threshold, we calculated the frequency of reads at each position that agreed with the mutation (derived) allele. As the sequencing depth was limited and *B. fragilis* represented only ∼5% of reads on average (**Extended Data Fig. 9a**), not every SNP was covered at every time point. For each SNP, we visualized its dynamics by using time points with non-zero read counts and smoothing the trajectory using the Savitzky-Golay method with a span of 25 and degree of 0 (**Fig. 4b**).

To plot a schematic of the population dynamics of different sublineages (**Fig. 4c**), we averaged frequencies of SNPs that were shared by a particular sublineage to estimate the relative abundance of this sublineage. To fill the time points where no stool community was sampled, we generated a continuous relative abundance trajectory for each sublineage using Fourier curve fitting (Matlab model fourier8). To visualize parent and child sublineages separately, we subtracted the relative abundance of a parent sublineage by the sum of relative abundances of its child sublineages. When the combined relative abundance of child sublineages exceeded that of their parent sublineage, we set the frequency of the parent sublineage to 0. After Day 180, we manually set the frequency of the SL1 parent genotype to zero, and reduced discontinuities caused by this assignment by an additional Fourier curve fitting step (Matlab parameter: fourier8). The imputed relative frequencies were then renormalized so that they sum up to 1.

We also examined L03’s dynamics during colonization using 74 metagenomes collected over 144 days (**Extended Data Fig. 9c-f**). The same methods were used as described above, with the exception that mutations in ≧3 isolates were able to be tracked, owing to the higher relative abundance of *B. fragilis* in Subject 03. This schematic shows an expansion of a SNP and SNPs that decrease over time.

### Data availability

Data is in the process of being uploaded to public servers. FASTQ files for the 602 *B. fragilis* isolates, with adaptors removed and filtered for quality, will be uploaded to the SRA. BAM files of the 352 metagenomes aligned to *B. fragilis* lineage assemblies will also be available on the SRA. Lineage assemblies with annotations will be uploaded to NCBI.

### Code availability

Commented custom MATLAB code will be uploaded to Github prior to publication.

---

**Extended Data 1:**
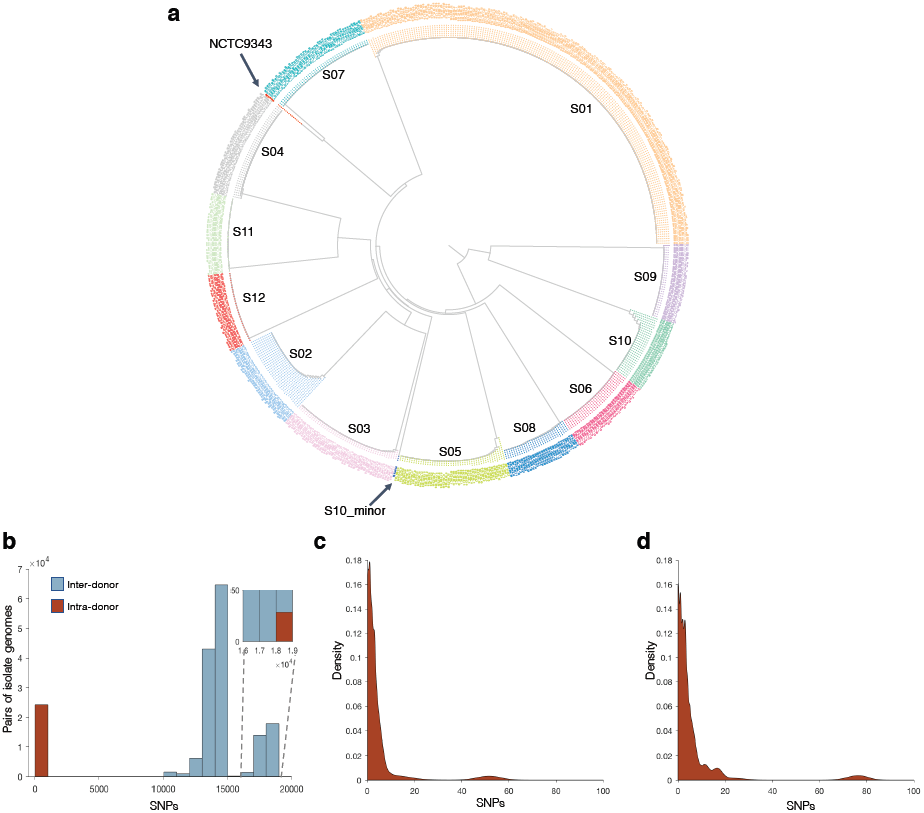
Inter-subject and intra-subject mutational distances between pairs of isolates suggest that each individual subject has a single dominant *B. fragilis* lineage. (**A**) Neighbor-joining tree of all isolates from 12 subjects, with nodes annotated with isolate IDs (Methods). Isolates from the same subject, except one isolate from S10, always closely clustered. Each cluster was uniquely colored and labeled by the subject ID. (**b**) Histogram of the mutational distances between all pairs of isolates. Inter-subject pairs are shown in blue, while intra-subject pairs are in red. The bin size is 1000 SNPs. Twenty-nine intra-subject pairs are >18000 SNPs apart and emerged from one isolate from Subject 10 that was from a minor lineage. (**c**) Excluding this minor lineage, all intra-subject mutational distances were <100 SNPs. The probability distribution of intra-subject mutational distances, averaging across 12 subjects, is shown. (**d**) The same analysis as in panel (**c**), but intra-lineage SNPs were identified using lineage assemblies as references. We observed a shift rightwards, suggesting assembly-based approaches identify more mutations.

**Extended Data 2:**
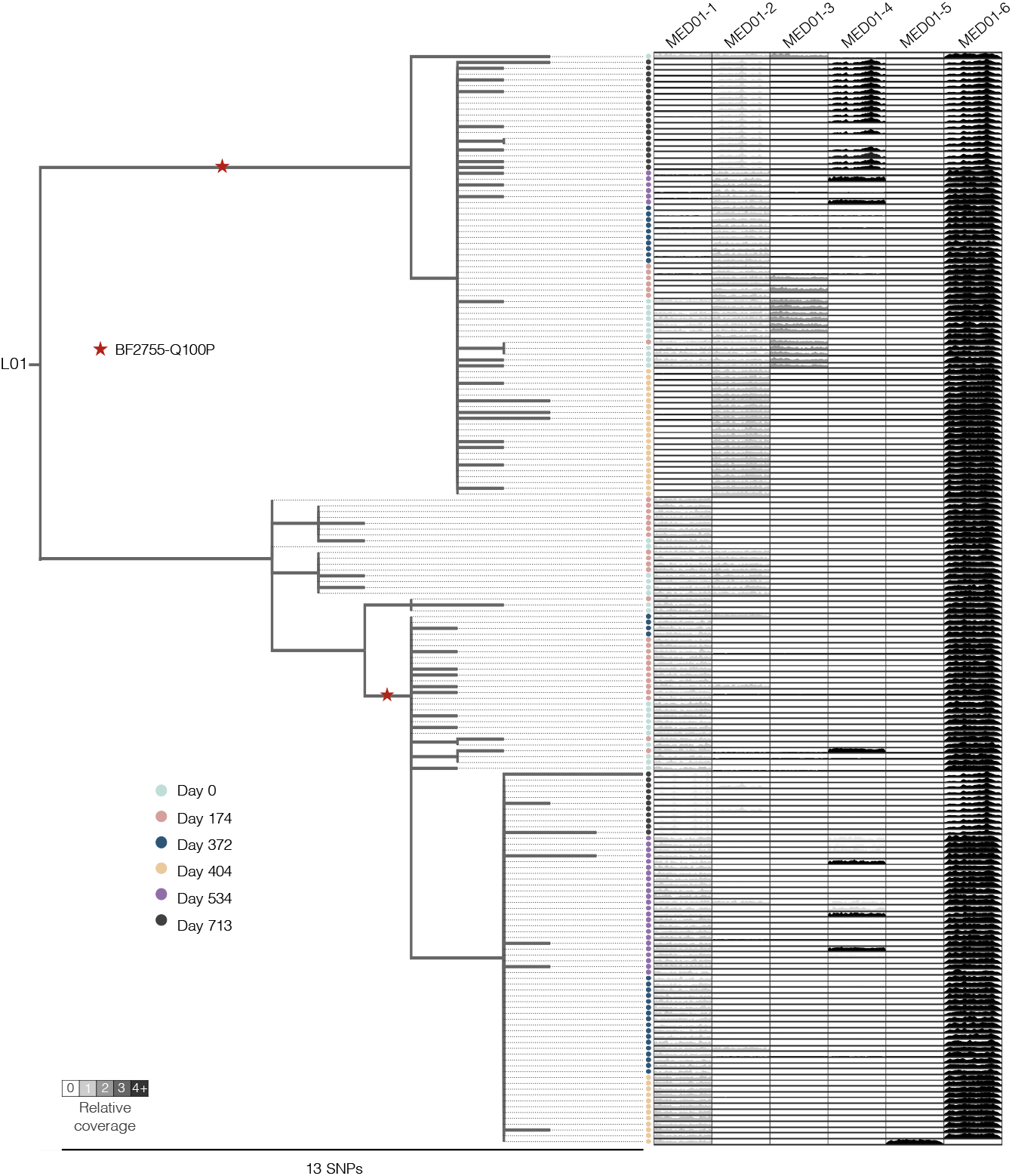
Within-person *B. fragilis* evolution in L01. The phylogeny for isolates from L01 is shown. Colored circles represent isolates from samples collected at the indicated dates. For each isolate, the relative coverage across identified MEDs is shown. Shading of MED regions reflects the average relative coverage of the MED in that isolate. Red stars indicate when the same nucleotide mutation emerged multiple times within the same lineage (inferred via parsimony, Methods). Isolates from Day 713 have different patterns of relative coverage across the MEDs because genomic libraries for these isolates were prepared differently (Method). More details on the exact mutations and MEDs found are in **Supplementary Tables 7** and **Supplementary Table 3**.

**Extended Data 3:**
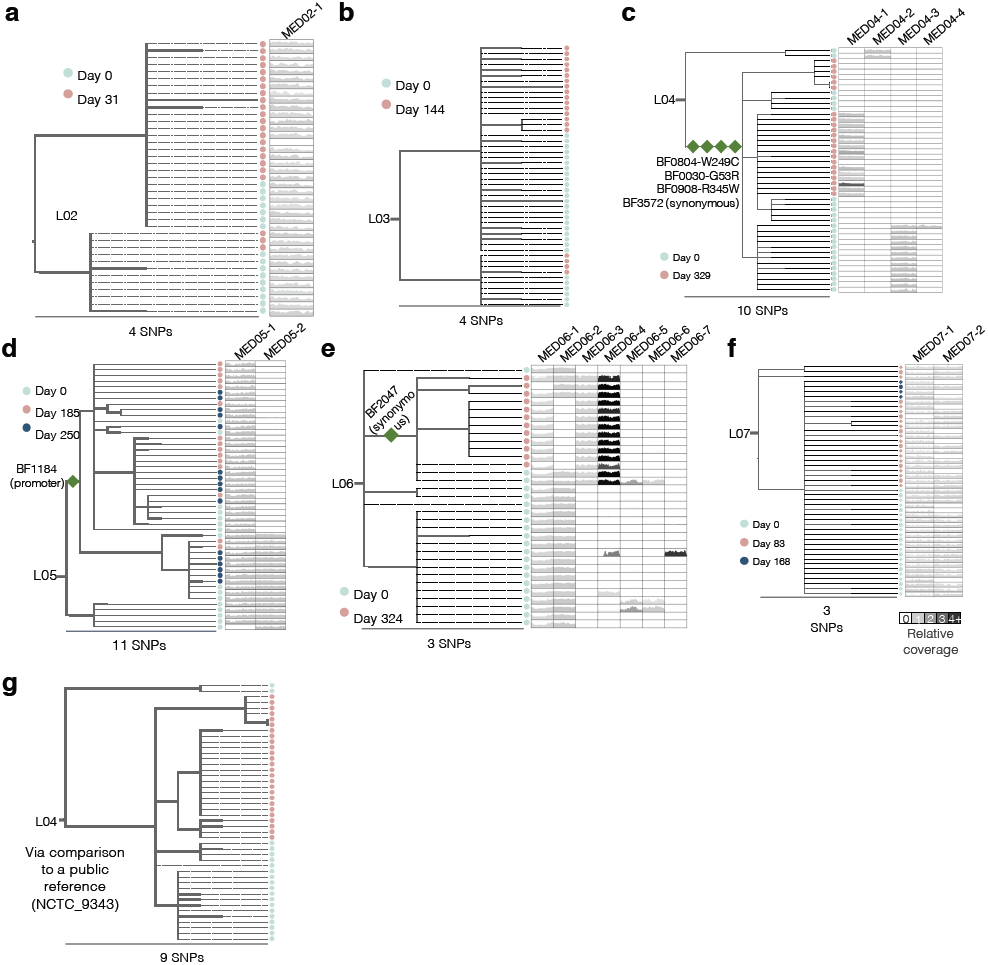
Within-person *B. fragilis* evolution in L02-L07. The phylogeny for isolates from L02 to L7, respectively. Colored circles represent isolates from samples collected at the indicated dates. For each isolate, the relative coverage across identified MEDs is shown. Shading of MED regions reflects the average relative coverage of the MED in that isolate. Dark green diamonds indicate SNPs associated with sweeps and are labeled with gene ID and type of mutation. In (**f**), the SNP that was shared by all isolates from the latest time point (dark blue) was not included as a sweep because it might be an artifact of undersampling at the later timepoint (**Extended Data Fig. 6g**). (**g**) Phylogeny for isolates from L04, via comparison to NCTC_9343 reference. No MEDs were identified using this public reference. More details on the exact mutations and MEDs identified from these lineages are in **Supplementary Tables 8-13** and **Supplementary Table 3**.

**Extended Data 4:**
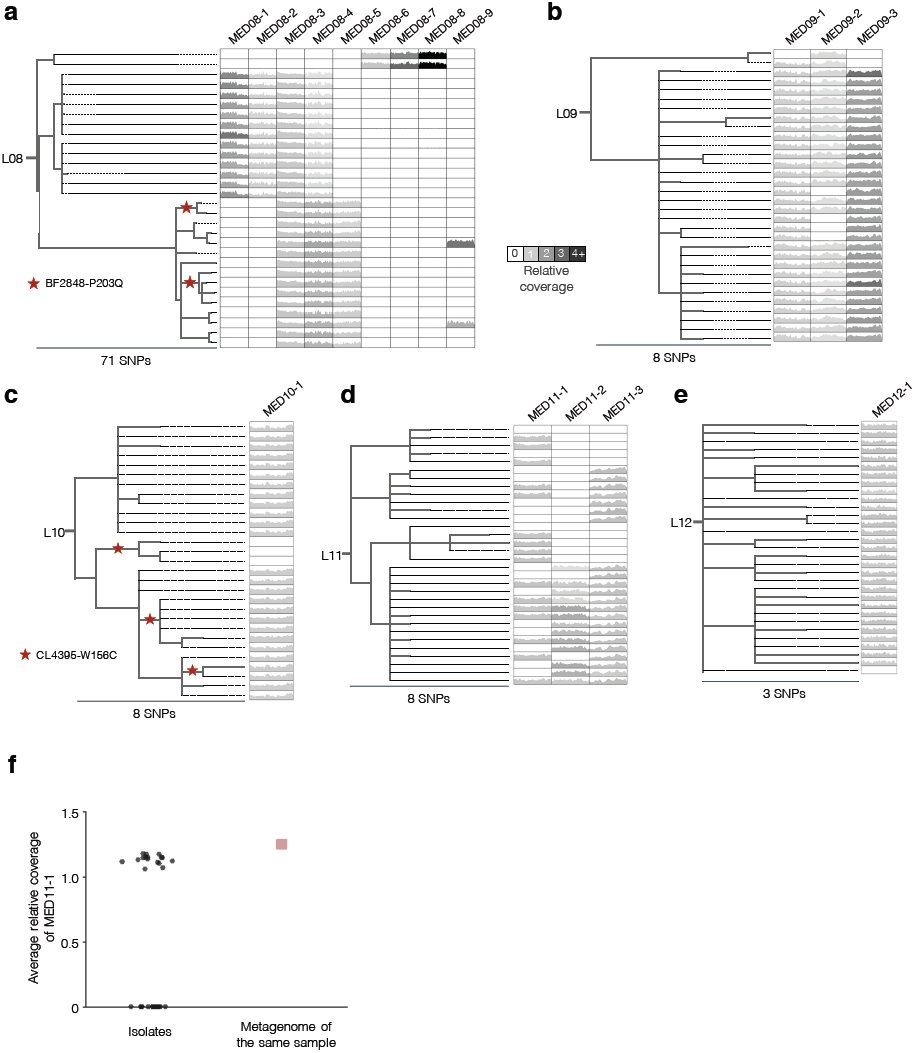
Within-person *B. fragilis* evolution in L08-L12. The phylogeny for isolates from L08 to L12, respectively. All lineages were sampled once. For each isolate, the relative coverage across identified MEDs is shown. Shading of MED regions reflects the average relative coverage pattern of the MED in that isolate. Red stars indicate when the same nucleotide mutation emerged multiple times within the same lineage (inferred via parsimony, Methods). (**d**) The presence/absence pattern of MED11-1 suggests many loss events on the phylogeny. (**f**) Notably, the relative coverage in the metagenome from the same sample is comparable to the relative coverage in individual isolates with the MED. This suggests that the MED may have been present in all cells and subsequently lost many times after stool collection from Subject 11, possibly during the culturing period. More details on the exact mutations and MEDs identified from these lineages are in **Supplementary Tables 14-18** and **Supplementary Table 3.**

**Extended Data 5:**
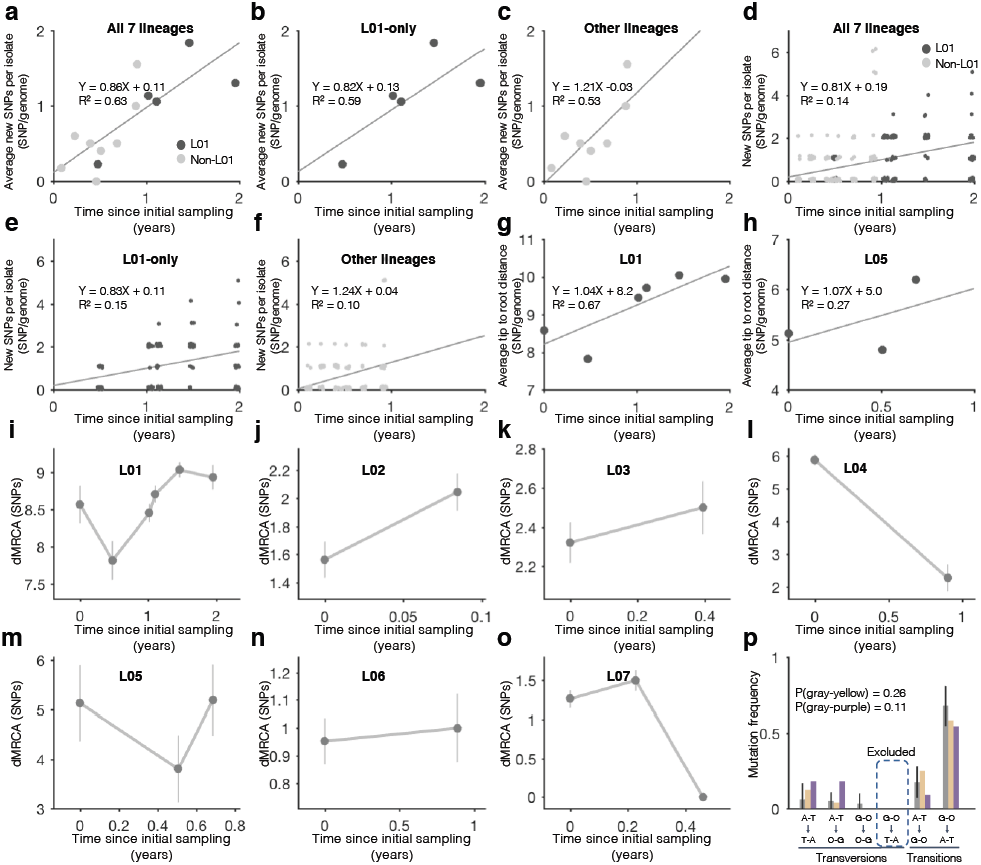
Further analysis on molecular clock, mutation spectrum and dMRCA changes over time. We used the number of new SNPs from isolates of later time points to compute the molecular clock (Methods, panel **a-f**). The molecular clock was calculated using the average number of new SNPs per isolate from the same time point (**a**) or new SNPs from each isolate (**d**), using all lineages with longitudinal samples. We performed the same calculations using only L01 (**b** and **e**) and all lineages (L02-L07, **c** and **f**). Each estimation gives similar values, but levels of R^2^ are much lower when considering isolates alone, due to variation in dMRCA between isolates taken at the same point. (**g**) We also calculated the molecular clock using the average tip-to-root distances of isolates from different time points from L01 (**g**) and L05 (**h**). This approach gives comparable estimates to the clock based on new SNPs. (**i-o**) For each time point of each subject that had longitudinal samples, we inferred the most recent common ancestor (MRCA) of just those isolates, and calculated dMRCA of the isolate population relative to that ancestor. Dots represent mean of dMRCA of each time points, error bars represent standard deviation of the mean. In 2 subjects (panels **l, m**), dMRCA decreases between time points were associated with SNPs fixed between samplings (**Extended Data Fig. 2c, d**). The decrease of dMRCA in L07 (**p**) was possibly an artifact due to an undersampling of the last time point (**Extended Data Fig. 6g**). Between time points 1 and 2 in L01, dMRCA also decreased, and this decrease was due to changes in relative abundances of sublineages with different distances to the (same) MRCA (**Extended Data Fig. 2**). (**n)** A sweep in L06 (**Extended Data Fig. 2e**) was not associated with a decrease in dMRCA. (**p**) Mutation spectrum analysis indicates no difference of hypermutator lineage from other lineages when GC to TA mutations are excluded (**Methods, Fig 1f**).

**Extended Data 6:**
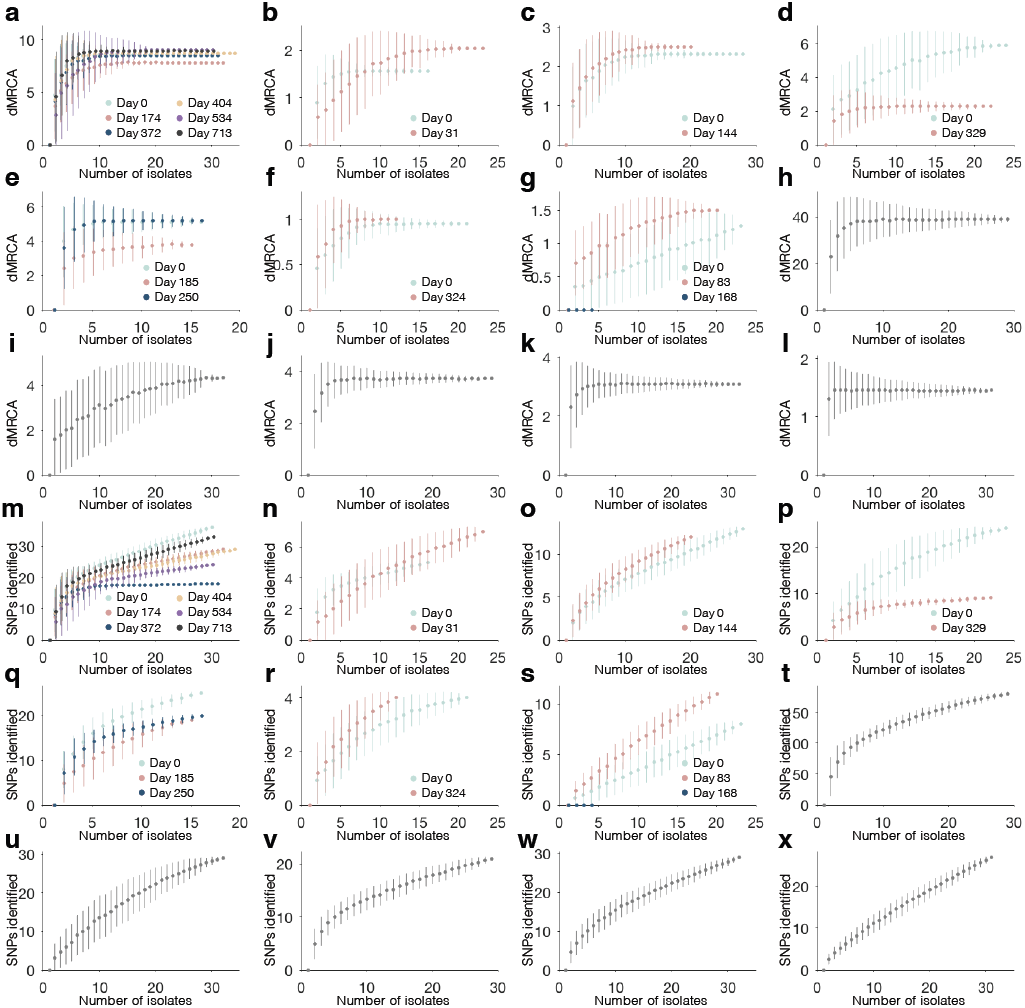
Collector curves suggest sufficient sampling for dMRCA, yet numbers of SNPs identified depends on number of isolates collected. (**A-L**) For each lineage and time point, we created a collector curve for dMRCA (one curve if the lineage was sampled once). For an isolate population from a particular time point, we subsampled the population to x isolates (0<x<n, n = total number of isolates at the time point), reconstructed the MRCA, and recomputed dMRCA. For each x, we simulated 100 subsamples and computed the mean (dots) and standard deviation (bars) for the simulation results. dMRCA was undersaturated only in 2 time points from L07 (0 and 168 Days). (**m-x**) For each lineage and time point, we created a collector curve for the number of SNPs identified (one curve if the lineage was sampled once). For an isolate population from a particular time point, we subsampled the population to x isolates (0<x<n, n = total number of isolates at the time point), and recomputed the number of SNPs identified. For each x, we simulated 100 subsamples and computed the mean (dots) and standard deviation (bars) for the simulation results.

**Extended Data 7:**
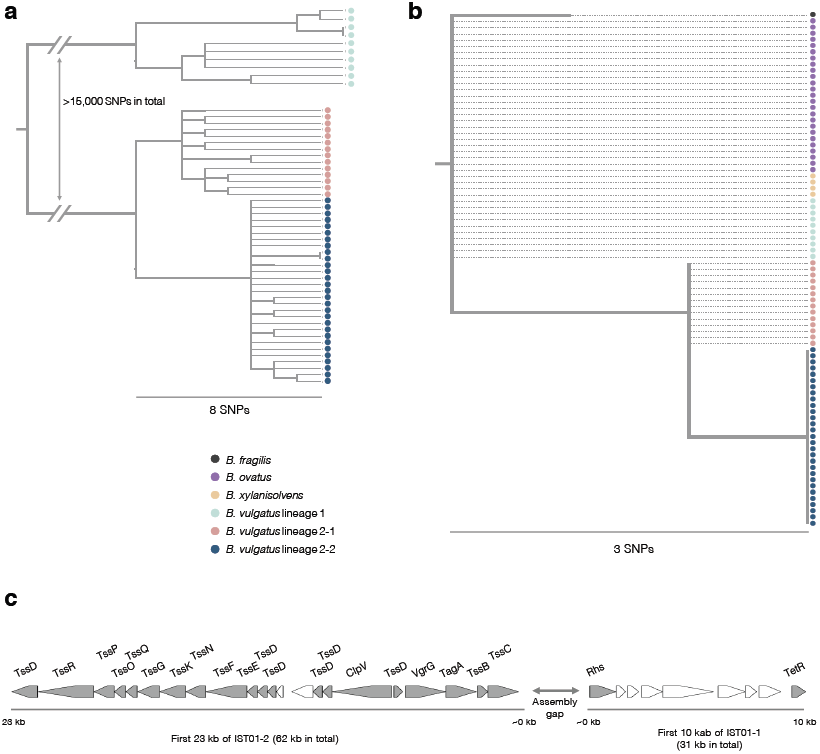
Transfer of a putative integrative conjugative element with type 6 secretion system across *Bacteroides* species within Subject 01. Analysis of the integrative conjugative element (ICE) found to be transferred in L01, identified from two candidate interspecies transfer regions (IST01-1 and IST01-2, Methods). (**a**) A phylogeny was constructed for all *B. vulgatus* isolates cultured from Subject 01, using a publicly available reference genome (GCF_000012825.1) and the same parameters and methods for *B. fragilis* SNP identification and evolutionary inference. We identified two *B. vulgatus* lineages that were separated by >15,000 SNPs. Within *B. vulgatus* lineage 2, we observed two sublineages. (**b**) A phylogeny was built using reads aligned to the ICE from all isolates of 4 *Bacteroides species* from Subject 01 (**Fig. 2d**). The sequences of IST01-1 and IST01-2 in the L01 assembly were used as the reference and the same methods were used as for *B. fragilis* SNP evolutionary inference. Among the 4 SNPs identified, we found 2 SNP locations whose 200-bp flanking sequence had matches in NCBI with >85% similarity, and we used these alleles as outgroups to root the tree. For the remaining 2 SNP locations, we assigned ancestral alleles that minimized the variance of dMRCA of all isolates. Colors represent isolates from the same phylogenetic group. The consensus ICE sequence in the L01 *B. fragilis* genome is represented by a single circle (black). We note that three SNPs were identified within *B. fragilis* L01, each in a single isolate. (**c**) ORF map of the type 6 secretion system of architecture 2 (T6SS-GA2) carried on this ICE. We aligned the ORFs from IST-01 and IST-02 to an annotated T6SS-GA2 from *Parabacteroides distasonis* CL03T12C09 (accession: JH976496.1). The first 10 kb of IST01-1 and the first 23 kb of IST01-2 had ORFs that are homologous to this T6SS-GA2. Gray pentagons represent conserved genes for T6SS-GA2^34^.

**Extended Data 8:**
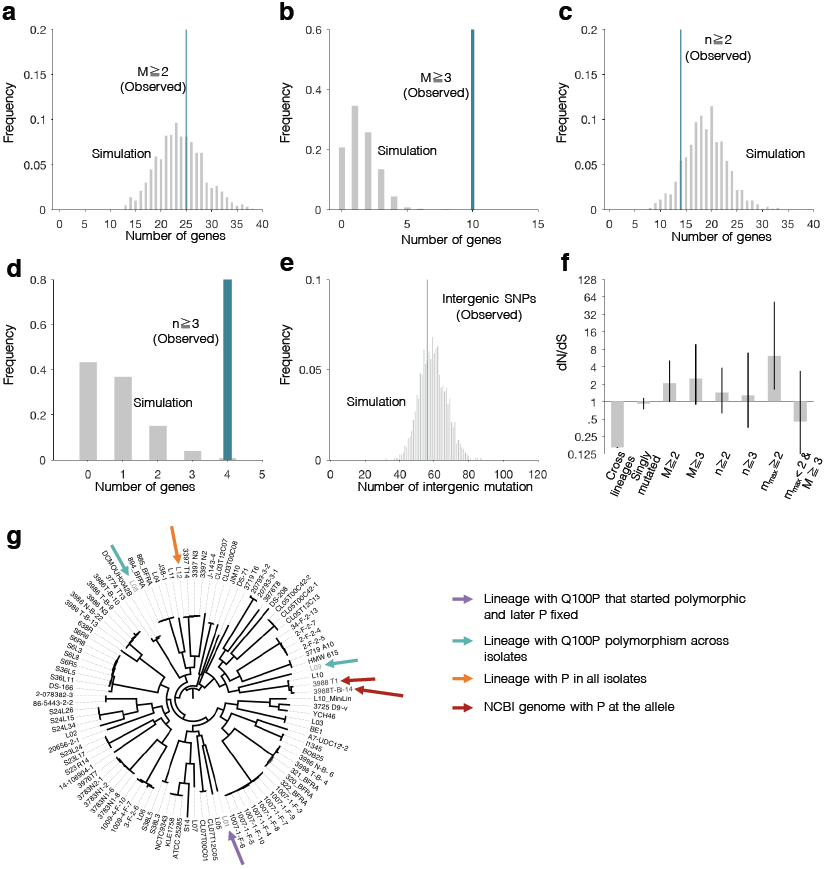
Search for parallel evolution across lineages did not yield additional genes under selection, and parallel evolution of Q100P in BF2755 across and within lineages. We searched for genes mutated multiple times across lineages, counting the number of total SNPs obtained in each gene (M), the number of lineages a gene was mutated in (n), and the maximum number of mutation a given gene was mutated in any lineage (m_max_). Simulations were performed as described in the Methods. (**a**) A search with the criteria of M≧2 yielded results consistent with a null model. (**b**) When this threshold was increased to M≧3, 11 genes were observed. Interestingly, 9 of these genes were already discovered with the criteria used in the main text, m_max_≧2. The 2 genes that are newly discovered with this metric (m_max_<2 & M≧3) do not show a signal for positive selection (**f**). (**c-d**) Similar results were obtained for the metric n, with the only 2 new genes discovered being identical to the analysis in (**a-b**). Further, dN/dS of genes discovered with the n metric did not show a significant signal for adaptive evolution (**f**). (**e**) The number of intergenic mutations is consistent with a null model. (**f**) dN/dS calculated across groups of genes defined with various metrics for parallel evolution. Together, these results are consistent with the evidence of person-specific selection forces found in the main text, and suggest that when a selection pressures is shared across subjects, it can usually be detected from just studying a single subject. (**g**) We built a phylogeny of genomes from 12 major lineages, 1 minor lineage from L10 and 88 references from NCBI. We clustered coding sequences from these 101 genomes with 95% similarity using CD-HIT and identified 277 genes present in all genomes. The number of shared genes is an underestimate, as the available NCBI genome references contain assemblies with varying quality. We performed a multiple sequence alignment for each shared gene using MAFFT v7.310^66^ and concatenated the alignment files. A phylogenetic tree was constructed using the GTRGAMMAI model from RaxML v8.2.11 (parameters: -m GTRGAMMAI –p 12345 -# 20)^67^. For the 12 major lineages we investigated, two lineages had both isolates with a glutamine (Q) and isolates with a proline (P) at position 100 in the BF2755 protein (L08, L09). L01 started with two distinct Q100P mutations (**Extended Data Fig. 2**) and later on the mutant genotype (P) fixed in the population. All isolates from the L12 lineage had a P at this position, suggesting it had fixed prior to or during colonization. The remaining 8 lineages did not have a mutation at this position. For 88 publicly available genomes, we comapared their genomes to the DNA sequence of BF2755, and examined the position of this mutation. Two lineages, 3988 T1 and 3988-B-14, encode for a P at this position.

**Extended Data 9:**
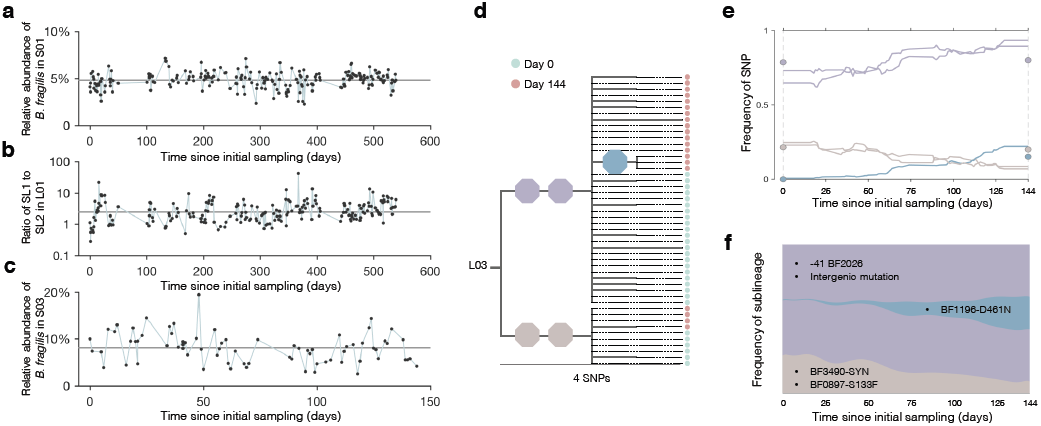
Evolutionary dynamics of L01 and L03. (**A**) For each metagenome from stool samples from Subject 01 (**Supplementary Table 6, Figure 4**), we calculated the percentage of metagenomic reads that aligned to the L01 genome assembly and plotted it against the time of sample collection. Reads potentially from other species (in regions with >5X median coverage) were excluded. This percentage estimates the relative abundance of *B. fragilis* in the stool community. The black line indicates the mean across samples. (**b**) For each sample, the ratio of SL1:SL2 was estimated using total number of reads aligned to alleles corresponding to either sublineage at the SNPs that separate them. Samples with fewer than 40 reads aligned to these SNP locations were excluded. The black line indicates the mean across samples. (**c**) The relative abundance of L03 *B. fragilis* inside Subject 03 was estimated in 74 metagenomes spanning 144 days, using the same method described in (**a**). (**d**) The phylogeny of isolates from L03. Branches with ≧3 isolates are labeled with colored octagons that represent individual SNPs. Circles represent individual isolates and are colored according to sampling date. (**e**) Frequencies of labeled SNPs over time in the *B. fragilis* population were inferred from 75 stool metagenomes (Methods). Colored circles represent SNP frequencies inferred from isolate genomes at particular time points. (**f**) The evolutionary history of sublineages during sampling was inferred (see Methods). Sublineages are defined by their signature SNPs, and labeled with the identity of SNPs and colored as in (**d**).

**Extended Data 10:**
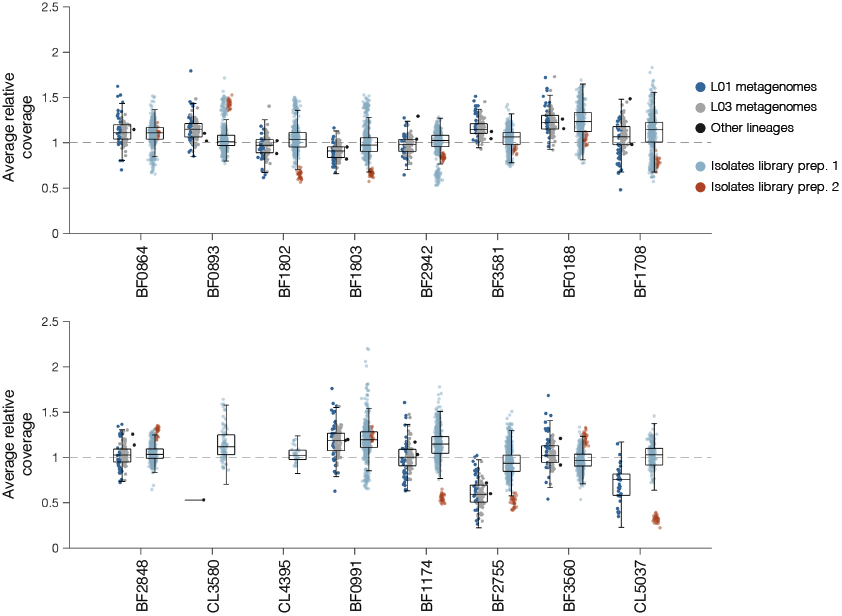
Coverage of the 16 genes under selection from metagenomes and isolate whole genomes. We recruited metagenomic reads to the assembled genomes and calculated the average relative coverage in the 16 genes under selection, if the gene was present in the lineage genome. Metagenomes with less than 10X coverage genomewide were excluded. The average relative coverages of these 16 genes were also computed for each of the isolates from all 12 lineages. For each gene, the relative coverage from metagenomes and isolates were grouped separately, and box and whiskers indicate their distributions. We note some minor differences between coverage in the metagenomes and isolates. We suspect these small differences are largely due to different library preparation methods, because isolate libraries prepared using different methods had similar differences in average relative coverage (Methods). Beyond this slight variation, we found no indication that any of these genes had abnormal coverage in the metagenomes, suggesting they are not shared with abundant species in the same microbiomes.

## Supplementary Tables

There are 19 Supplementary Tables uploaded in a single .xlsx file.

Supplementary Table 1: Subject information and per-lineage statistics

Supplementary Table 2: Stool samples used for culturing single-colony isolates

Supplementary Table 3: Mobile element difference (MED) information

Supplementary Table 4: Candidate inter-species transfers

Supplementary Table 5: Genes under selection *in vivo*

Supplementary Table 6: Stool samples used for metagenomic sequencing and alignment results

Supplementary Table 7: *de novo* SNPs within L01

Supplementary Table 8: *de novo* SNPs within L02

Supplementary Table 9: *de novo* SNPs within L03

Supplementary Table 10: *de novo* SNPs within L04

Supplementary Table 11: *de novo* SNPs within L05

Supplementary Table 12: *de novo* SNPs within L06

Supplementary Table 13: *de novo* SNPs within L07

Supplementary Table 14: *de novo* SNPs within L08

Supplementary Table 15: *de novo* SNPs within L09

Supplementary Table 16: *de novo* SNPs within L10

Supplementary Table 17: *de novo* SNPs within L11

Supplementary Table 18: *de novo* SNPs within L12

Supplementary Table 19: Clustering of gene homologs from different lineages

## References

1. Faith, J. J. et al. The Long-Term Stability of the Human Gut Microbiota. Science (80-.). 341, 1237439–1237439 (2013).

2. Schloissnig, S. et al. Genomic variation landscape of the human gut microbiome. Nature 493, 45–50 (2012).

3. Zoetendal, E. G., Akkermans, A. D. & De Vos, W. M. Temperature gradient gel electrophoresis analysis of 16S rRNA from human fecal samples reveals stable and host-specific communities of active bacteria. Appl. Environ. Microbiol. 64, 3854–9 (1998).

4. Ghalayini, M. et al. ‘Evolution of a dominant natural isolate of Escherichia coli in the human gut over a year suggests a neutral evolution with reduced effective population size’. Appl. Environ. Microbiol. AEM.02377-17 (2018). doi:10.1128/AEM.02377-17

5. Lee, S. M. et al. Bacterial colonization factors control specificity and stability of the gut microbiota. Nature 501, 426–429 (2013).

6. Sender, R., Fuchs, S. & Milo, R. Revised Estimates for the Number of Human and Bacteria Cells in the Body. PLoS Biol. 14, 1–14 (2016).

7. Barrick, J. E. & Lenski, R. E. Genome dynamics during experimental evolution. Nat. Rev. Genet. 14, 827–39 (2013).

8. Nayfach, S. & Pollard, K. S. Average genome size estimation improves comparative metagenomics and sheds light on the functional ecology of the human microbiome. Genome Biol. 16, 51 (2015).

9. Korem, T. et al. Growth dynamics of gut microbiota in health and disease inferred from single metagenomic samples. Science (80-.). 349, 1101–1106 (2015).

10. Didelot, X., Walker, A. S., Peto, T. E., Crook, D. W. & Wilson, D. J. Within-host evolution of bacterial pathogens. Nat. Rev. Microbiol. 14, 150–162 (2016).

11. Lieberman, T. D. et al. Parallel bacterial evolution within multiple patients identifies candidate pathogenicity genes. Nat. Genet. 43, 1275–1280 (2011).

12. Marvig, R. L., Sommer, L. M., Molin, S. & Johansen, H. K. Convergent evolution and adaptation of Pseudomonas aeruginosa within patients with cystic fibrosis. Nat. Genet. 47, 57–64 (2015).

13. Feliziani, S. et al. Coexistence and Within-Host Evolution of Diversified Lineages of Hypermutable Pseudomonas aeruginosa in Long-term Cystic Fibrosis Infections. PLoS Genet. 10, (2014).

14. Kennemann, L. et al. Helicobacter pylori genome evolution during human infection. Proc. Natl. Acad. Sci. 108, 5033–5038 (2011).

15. Golubchik, T. et al. Within-Host Evolution of Staphylococcus aureus during Asymptomatic Carriage. PLoS One 8, 1–14 (2013).

16. Groussin, M. et al. Unraveling the processes shaping mammalian gut microbiomes over evolutionary time. Nat. Commun. 8, 14319 (2017).

17. Goodrich, J. K. et al. Genetic Determinants of the Gut Microbiome in UK Twins. Cell Host Microbe 19, 731–43 (2016).

18. He, M. et al. Evolutionary dynamics of Clostridium difficile over short and long time scales. Proc. Natl. Acad. Sci. 107, 7527–7532 (2010).

19. Huttenhower, C. et al. Structure, function and diversity of the healthy human microbiome. Nature 486, 207–214 (2012).

20. Chattopadhyay, S. et al. High frequency of hotspot mutations in core genes of Escherichia coli due to short-term positive selection. Proc. Natl. Acad. Sci. U. S. A. 106, 12412–12417 (2009).

21. Barroso-Batista, J., Demengeot, J. & Gordo, I. Adaptive immunity increases the pace and predictability of evolutionary change in commensal gut bacteria. Nat. Commun. 6, 8945 (2015).

22. Garud, N. R., Good, B. H., Hallatschek, O. & Pollard, K. S. Evolutionary dynamics of bacteria in the gut microbiome within and across hosts. Doi.Org 210955 (2017). doi:10.1101/210955

23. Lieberman, T. D. et al. Genetic variation of a bacterial pathogen within individuals with cystic fibrosis provides a record of selective pressures. Nat Genet 46, 82–87 (2014).

24. Verster, A. J. et al. The Landscape of Type VI Secretion across Human Gut Microbiomes Reveals Its Role in Community Composition. Cell Host Microbe 22, 411–419.e4 (2017).

25. Chen, L. et al. VFDB: a reference database for bacterial virulence factors. Nucleic Acids Res. 33, D325–D328 (2004).

26. Yassour, M. et al. Natural history of the infant gut microbiome and impact of antibiotic treatment on bacterial strain diversity and stability. Sci. Transl. Med. 8, 343ra81–343ra81 (2016).

27. David, L. A. et al. Host lifestyle affects human microbiota on daily timescales. Genome Biol. 15, R89 (2014).

28. Lozupone, C. A., Stombaugh, J. I., Gordon, J. I., Jansson, J. K. & Knight, R. Diversity, stability and resilience of the human gut microbiota. Nature 489, 220–230 (2012).

29. Giraud, A. Costs and Benefits of High Mutation Rates: Adaptive Evolution of Bacteria in the Mouse Gut. Science (80-.). 291, 2606–2608 (2001).

30. Chu, N. D. et al. A Mobile Element in mutS Drives Hypermutation in a Marine Vibrio. MBio 8, e02045–16 (2017).

31. Jolivet-Gougeon, A. et al. Bacterial hypermutation: clinical implications. J. Med. Microbiol. 60, 563–573 (2011).

32. Marvig, R. L., Johansen, H. K., Molin, S. & Jelsbak, L. Genome Analysis of a Transmissible Lineage of Pseudomonas aeruginosa Reveals Pathoadaptive Mutations and Distinct Evolutionary Paths of Hypermutators. PLoS Genet. 9, (2013).

33. Matic, I. Highly Variable Mutation Rates in Commensal and Pathogenic Escherichia coli. Science (80-.). 277, 1833–1834 (1997).

34. Coyne, M. J., Roelofs, K. G. & Comstock, L. E. Type VI secretion systems of human gut Bacteroidales segregate into three genetic architectures, two of which are contained on mobile genetic elements. BMC Genomics 17, 58 (2016).

35. Coyne, M. J. et al. Evidence of Extensive DNA Transfer between Bacteroidales Species within the Human Gut. MBio 5, e01305–14 (2014).

36. Cerdeno-Tarraga, A. M. Extensive DNA Inversions in the B. fragilis Genome Control Variable Gene Expression. Science (80-.). 307, 1463–1465 (2005).

37. Martens, E. C., Koropatkin, N. M., Smith, T. J. & Gordon, J. I. Complex glycan catabolism by the human gut microbiota: The bacteroidetes sus-like paradigm. J. Biol. Chem. 284, 24673–24677 (2009).

38. Glenwright, A. J. et al. Structural basis for nutrient acquisition by dominant members of the human gut microbiota. Nature 541, 407–411 (2017).

39. Woude, M. W. Van Der & Bäumler, A. J. Phase and Antigenic Variation in Bacteria Phase and Antigenic Variation in Bacteria. Clin. Microbiol. Rev. 17, 581–611 (2004).

40. Burt, S., Meldrum, S., Woods, D. R. & Jones, D. T. Colonial variation, capsule formation, and bacteriophage resistance in Bacteroides thetaiotaomicron. Appl. Environ. Microbiol. 35, 439–443 (1978).

41. Messer, P. W., Ellner, S. P. & Hairston, N. G. Can Population Genetics Adapt to Rapid Evolution? Trends Genet. 32, 408–418 (2016).

42. Bell, G. Fluctuating selection: the perpetual renewal of adaptation in variable environments. Philos. Trans. R. Soc. Lond. B. Biol. Sci. 365, 87–97 (2010).

43. Krinos, C. M. et al. Extensive surface diversity of a commensal microorganism by multiple DNA inversions. Nature 414, 555–558 (2001).

44. Porter, N. T., Canales, P., Peterson, D. A. & Martens, E. C. A Subset of Polysaccharide Capsules in the Human Symbiont Bacteroides thetaiotaomicron Promote Increased Competitive Fitness in the Mouse Gut. Cell Host Microbe 22, 494–506 (2017).

45. Kuwahara, T. et al. Genomic analysis of Bacteroides fragilis reveals extensive DNA inversions regulating cell surface adaptation. Proc. Natl. Acad. Sci. 101, 14919–14924 (2004).

46. Rocabert, C., Knibbe, C., Consuegra, J., Schneider, D. & Beslon, G. Beware batch culture: Seasonality and niche construction predicted to favor bacterial adaptive diversification. PLOS Comput. Biol. 13, e1005459 (2017).

47. Chung, H. et al. Global and local selection acting on the pathogen Stenotrophomonas maltophilia in the human lung. Nat. Commun. 8, 14078 (2017).

48. Good, B. H., McDonald, M. J., Barrick, J. E., Lenski, R. E. & Desai, M. M. The dynamics of molecular evolution over 60,000 generations. Nature (2017). doi:10.1038/nature24287

49. Plucain, J. et al. Epistasis and allele specificity in the emergence of a stable polymorphism in Escherichia coli. Science (80-.). 1242862 (2014).

50. Smith, E. E. et al. Genetic adaptation by Pseudomonas aeruginosa to the airways of cystic fibrosis patients. Proc. Natl. Acad. Sci. U. S. A. 103, 8487–92 (2006).

## Methods references

51. Baym, M. et al. Inexpensive Multiplexed Library Preparation for Megabase-Sized Genomes. PLoS One 10, e0128036 (2015).

52. Martin, M. Cutadapt removes adapter sequences from high-throughput sequencing reads. EMBnet J 17, 10–12 (2011).

53. Joshi, N. A. & Fass, J. N. Sickle: A sliding-window, adaptive, quality-based trimming tool for FastQ files (Version 1.33)[Software]. (2011).

54. Li, H. et al. The Sequence Alignment/Map format and SAMtools. Bioinformatics 25, 2078–2079 (2009).

55. Bankevich, A. et al. SPAdes: A New Genome Assembly Algorithm and Its Applications to Single-Cell Sequencing. J. Comput. Biol. 19, 455–477 (2012).

56. Seemann, T. Prokka: Rapid prokaryotic genome annotation. Bioinformatics 30, 2068–2069 (2014).

57. Plotree, D. & Plotgram, D. PHYLIP-phylogeny inference package (version 3.2). cladistics 5, 6 (1989).

58. Fu, L., Niu, B., Zhu, Z., Wu, S. & Li, W. CD-HIT: accelerated for clustering the next-generation sequencing data. Bioinformatics 28, 3150–3152 (2012).

59. Price, M. N. & Arkin, A. P. PaperBLAST: Text Mining Papers for Information about Homologs. mSystems 2, e00039–17 (2017).

60. Ashkenazy, H., Erez, E., Martz, E., Pupko, T. & Ben-Tal, N. ConSurf 2010: Calculating evolutionary conservation in sequence and structure of proteins and nucleic acids. Nucleic Acids Res. 38, 529–533 (2010).

61. Joint Center for Structural Genomics. Crystal structure of a SusD superfamily protein (BF1802) from Bacteroides fragilis NCTC 9343 at 1.90 A resolution. The Protein Data Bank (2010). doi:10.2210/pdb3nqp/pdb

62. McWilliam, H. et al. Analysis Tool Web Services from the EMBL-EBI. Nucleic Acids Res. 41, W597–W600 (2013).

63. Schrödinger, LLC. The {PyMOL} Molecular Graphics System, Version∼1.8. (2015).

64. Krissinel, E. & Henrick, K. Inference of Macromolecular Assemblies from Crystalline State. J. Mol. Biol. 372, 774–797 (2007).

65. Yu, C.-S., Chen, Y.-C., Lu, C.-H. & Hwang, J.-K. Prediction of protein subcellular localization. Proteins Struct. Funct. Bioinforma. 64, 643–651 (2006).

## Extended Data references

Yamada, K. D., Tomii, K. & Katoh, K. Application of the MAFFT sequence alignment program to large data - Reexamination of the usefulness of chained guide trees. Bioinformatics 32, 3246–3251 (2016).

Stamatakis, A. RAxML-VI-HPC: Maximum likelihood-based phylogenetic analyses with thousands of taxa and mixed models. Bioinformatics 22, 2688–2690 (2006).

